# Human anti-ACE2 monoclonal antibodies as pan-sarbecovirus prophylactic agents

**DOI:** 10.1101/2022.08.24.505169

**Authors:** Fengwen Zhang, Jesse Jenkins, Renan V.H. de Carvalho, Sandra Nakandakari-Higa, Teresia Chen, Morgan E. Abernathy, Elisabeth Nyakatura, David Andrew, Irina Lebedeva, Ivo C Lorenz, H.-Heinrich Hoffmann, Charles M. Rice, Gabriel D. Victora, Christopher O. Barnes, Theodora Haziioannou, Paul D. Bieniasz

**Affiliations:** Laboratory of Retrovirology, The Rockefeller University, New York, NY 10065; Laboratory of Lymphocyte Dynamics, The Rockefeller University, New York, NY 10065; Department of Biology, Stanford University, Stanford, CA 94305; Tri-institutional Therapeutics Discovery Institute, New York, NY 10021; Laboratory of Virology and Infectious Disease, The Rockefeller University, New York, NY 10065; Chan Zuckerberg Biohub, San Francisco, CA 94158; Howard Hughes Medical Institute, The Rockefeller University, New York, NY 10065

## Abstract

Human monoclonal antibodies from convalescent individuals that target the SARS-CoV-2 spike protein have been deployed as therapeutics against SARS-CoV-2. However, nearly all of these antibodies have been rendered obsolete by SARS-CoV-2 variants that evolved to resist similar, naturally occurring antibodies. Here, we describe the development of human monoclonal antibodies that bind the ACE2 receptor rather than the viral spike protein. These antibodies block infection by all ACE2 binding sarbecoviruses, including emergent SARS-CoV-2 variants. Structural and biochemical analyses revealed that the antibodies target an ACE2 epitope that engages SARS-CoV-2 spike. Importantly, the antibodies do not inhibit ACE2 enzymatic activity, nor do they induce ACE depletion from cell surfaces. The antibodies exhibit favorable pharmacology and protect human ACE2 knock-in mice against SARS-CoV-2 infection. Such antibodies should be useful prophylactic and treatment agents against any current and future SARS-CoV-2 variants, as well as ACE2-binding sarbecoviruses that might emerge as future pandemic threats.

## Introduction

Human monoclonal antibodies (mAbs) that target self or foreign antigens are a rich source of therapeutic molecules that can be administered as recombinantly produced proteins (Buss et al., 2012). In the context of viral infection, mAbs can confer a state of passive immunity in prophylactic or therapeutic approaches (Corti et al., 2021). Many groups, including our own, have developed human mAbs that target the SARS-CoV-2 spike protein as candidate therapeutic/preventive agents (Corti *et al*., 2021; 2022; Levin et al., 2022; Robbiani et al., 2020; Weinreich et al., 2021; Yang et al., 2020). However, the therapeutic or prophylactic use of spike-targeting antibodies against SARS-CoV-2, and viruses in general, has two key conceptual drawbacks. First, the mAbs are generated by human immune systems, and the most potent of them are identical to, or very similar to, potent neutralizing antibodies commonly elicited by infection or vaccination (Barnes et al., 2020; Corti *et al*., 2021; Wang et al., 2021). Consequently, as SARS-CoV-2 has replicated in human populations with increasing levels of infection or vaccination induced immunity, it has evolved to acquire resistance to therapeutic mAbs, even when these mAbs have not been widely used as therapeutics (Baum et al., 2020; Cao et al., 2022; Muecksch et al., 2021; Weisblum et al., 2020). Indeed, emergent SARS-CoV-2 variants, such as omicron and derivatives thereof, have rendered obsolete most SARS-CoV-2 mAb therapeutics generated from the antibody repertoires of individuals infected with prior SARS-CoV-2 variants (Cao *et al*., 2022; Schmidt et al., 2021a).

A second drawback of sarbecovirus spike-targeting mAbs that might be stockpiled in anticipation of future pandemics, is forecasting. Again, SARS-CoV-2 is an illustrative example; it is one of three recently emergent coronaviruses that are, to a large extent, antigenically distinct from each other (Sariol and Perlman, 2020). Without prior knowledge, it is near-impossible to pre-emptively generate spike targeting mAb therapeutics that would be predictably effective against an emergent virus, and variants thereof. Optimally, mAb therapeutics and prophylactics should be resilient to mutations selected during spread through human populations and ideally should be effective against entire classes of viruses (for example, the sarbecovirus subfamily).

Sarbecoviruses, including SARS-CoV, SARS-CoV-2 and SARS-related coronaviruses in bats and other mammals, use Angiotensin-converting enzyme-2 (ACE2) as the primary functional receptor for the spike glycoprotein (Letko et al., 2020; Li et al., 2003). In principle, antiviral breadth against sarbecoviruses and resilience to escape mutations can be achieved rather simply - by targeting mAbs to the human (h)ACE2 virus receptor rather than to sarbecovirus spike proteins. Resistance to such mAbs would require a profound change in how sarbecovirues interact with ACE2, or acquisition the ability to use a new receptor, a high genetic hurdle. While the use of mAbs targeting a self, cell-surface molecule might appear to carry greater risk of unwanted side effects than targeting viral spike proteins, there are precedents for the therapeutic deployment of receptor-blocking mAbs, particularly in situations when the viral or cellular ligands of such receptors are too variable to be blocked by a single mAb. For example, the HIV-1 receptor (CD4)-binding antibody Ibalizumab is approved as an HIV-1 therapeutic and is effective against a broad range of viral strains (Rizza et al., 2019). Other examples include treatments of conditions that involve diverse ligands such as interferonopathies; the 16 type-I interferons are too variable to be neutralized by a single antibody. However, anifrolumab, a mAb that inhibits the action of all type-I IFNs by binding the type-I IFN receptor is an approved therapeutic for systemic lupus erythematosus (Furie et al., 2017; Peng et al., 2015). Crucially, self, cell-surface targeting mAbs can be used safely in humans because Fc domains can be engineered to ablate cytotoxic effector functions (Tamm and Schmidt, 1997; Wang et al., 2018; Zalevsky et al., 2010).

Here, we generated a suite of hACE2 specific human mAbs that bind hACE2 with affinities in the low nM to pM range. These mAbs block infection by pseudotypes of all tested sarbecoviruses, with a potency that approaches that of the potent SARS-CoV-2 spike targeting therapeutic mAbs. A 3.3Å cryo-EM structure of one such mAb bound to hACE2 shows recognition of the α1 helix, leading to epitope competition with the spike receptor binding domain (RBD). Crucially, the mAbs do not inhibit ACE2 enzymatic activity or induce ACE2 internalization and have favorable pharmacology in hACE2 knock-in mice. Importantly, the mAbs provide protection against SARS-CoV-2 lung infection in hACE2 knock-in mice. The mAbs described herein thus have the potential to be used as prophylactic and treatment agents against current and any emergent SARS-CoV-2 variants as well as future sarbecovirus pandemic threats.

## Results

### Generation of hACE2-binding human monoclonal antibodies

Occlusion of the spike binding site on the human (h)ACE2 host receptors, using hACE2-binding mAbs has the potential to protect against infection. In a pilot experiment, BALB/c6 mice were immunized with recombinant hACE2 extracellular domain (1-740 aa). Sera collected at 35 days after immunization showed potent antiviral activity against SARS-CoV-2 pseudotyped viruses, with 50% inhibitory titers in the range 1860-6050 (Figure S1) and this result is consistent with findings that murine mAbs targeting hACE2 can inhibit sarbecovirus infection (Chen et al., 2021). This finding prompted us to generate human anti-hACE2 monoclonal mAbs through immunization of AlivaMab mice that generate chimeric human-mouse antibodies consisting of human Fab domains and a murine Fc domain. AlivaMab mice were immunized with recombinant hACE2 extracellular domains (1-740 aa) that were either monomeric and His-tagged or were fused to the Fc portion of human IgG1 to generate dimers, recapitulating the native hACE2 conformation. Hybridomas were generated from mice with sera exhibiting SARS-CoV-2 pseudotype infection-inhibiting activity and hybridoma supernatants were screened for hACE2 binding mAbs by enzyme-linked immunosorbent assay (ELISA) with plates coated with purified hACE2 (Figure 1A). Eighty-two hybridomas expressing hACE2-binding mAbs were identified. These anti-ACE2 mAb-containing hybridoma supernatants were tested for inhibition of pseudotyped virus infection of Huh-7.5 cells. Ten hybridomas secreting antibodies that potently inhibited SARS-CoV-2 pseudotyped virus infection (1C9H1, 2C12H3, 2F6A6, 2G7A1, 4A12A4, 05B04, 05D06, 05E10, 05G01, and 05H02, Figure 1A) were selected for subcloning. Thereafter, chimeric mAbs were purified from the hybridoma culture supernatants and antiviral activity confirmed in the SARS-CoV-2 pseudotyped virus inhibition assay. Additionally, the human Fab variable regions (VH and VL) were sequenced (Figure 1A, Table S1-S2). Four out of 5 mAbs derived from the KP AlivaMab mouse strain that generates human Kappa light chain (κ) containing antibodies, namely 05B04, 05D06, 05E10, and 05G01, shared similar or identical complementarity determining regions (CDRs) (Table S1, Table S2). In contrast, the mAbs from AV AlivaMab that generate both human Kappa (κ) and Lambda (λ) light chains were comparatively diverse and appeared to originate from distinct germline precursors (Table S1, Table S2).

**Figure 1.**
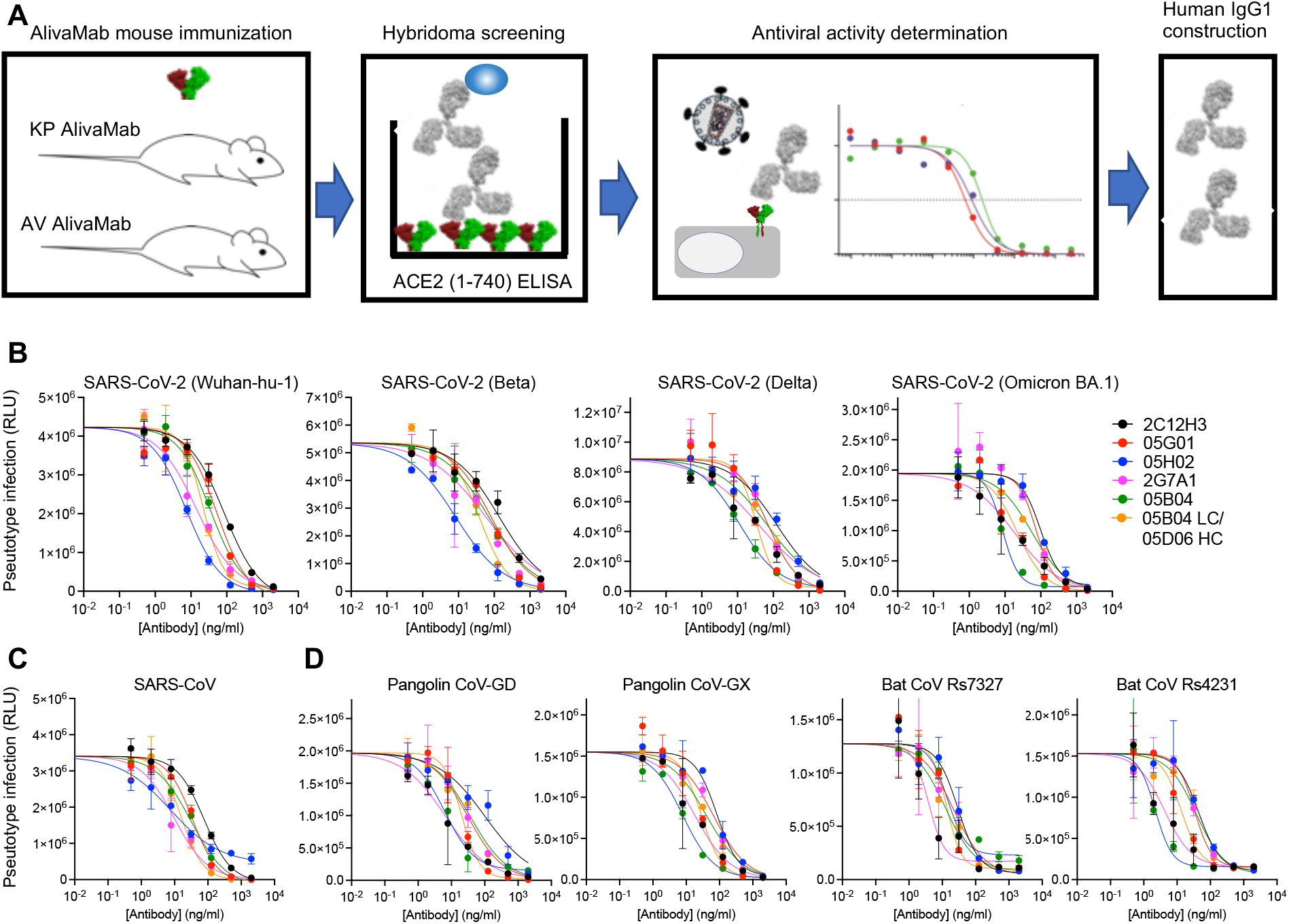
Discovery of human anti-human ACE2 antibodies that block sarbecovirus pseudotype infection. (A) Recombinant ACE2 extracellular domain (1-740 aa) as His-tagged (monomer) or Fc fused (dimer) proteins were injected into KP AlivaMab mice which generate human Kappa (κ) light chains or AV AlivaMab which generate both human Kappa (κ) and human Lambda (λ) light chains. ELISA screening yielded 82 reactive hybridoma clones. The supernatants of positive hybridoma clones were tested for inhibition of SARS-CoV-2 spike-pseudotyped virus infection and thereafter 10 human IgG1 antibodies were constructed. (B-D) Inhibition of HIV-1 based pseudotyped virus infection by anti-hACE2 mAbs. Six fully human anti-human ACE2 (hACE2) antibodies (2C12H3, 05G01, 05H02, 2G7A1, 05B04 and the hybrid antibody 05B04 LC/05D06 HC) were incubated with Huh-7.5 target cells and cells were infected with pseudotypes bearing spike proteins from SARS-CoV-2 variants (B) SARS-CoV (C) or Pangolin ans Bat sarbecoviruses viruses pseudotyped with the sarbecovirus spike proteins indicated. Infection was measured using NanoLuc luciferase assays. Mean and range of two independent titrations is plotted

### Human anti-hACE2 mAbs broadly inhibit sarbecovirus infection

To generate fully human anti-ACE2 mAbs, the variable domains from six of the chimeric human-mouse mAbs were inserted into a human IgG1 expression vector. The Fc domain was modified to include the mutations at L234/L235 (LALA) that abolish interacting with FcR-gamma (FcγR) and prevent cytotoxicity and at M428/N434 (LS) to enhance FcRn interaction and prolong half-life in humans (Tamm and Schmidt, 1997; Wang *et al*., 2018; Zalevsky *et al*., 2010). The fully human mAbs were generated by cotransfection with plasmids expressing corresponding heavy and light chains. Because the 05D06 human mAb generated only low yields, we expressed a hybrid mAb, 05B04LC/05D06HC, in which the light chain of 05B04 and the heavy chain of a clonally related mAb, 05D06 were cotransfected.

We tested the ability of the six mAbs to inhibit SARS-CoV-2 (Wuhan-hu-1) spike pseudotyped HIV-1 virus infection in Huh-7.5 target cells (Schmidt et al., 2020), which express endogenous ACE2 (Shang et al., 2021) and are permissive to SARS-CoV-2 infection (Zhu et al., 2020). Each of the mAbs tested were able to inhibit SARS-CoV-2 (Wuhan-hu-1) pseudotyped virus infection with IC_50_ values in the range of 7.5 ng/ml to 86 ng/ml (Figure 1B, Table 1). The most potent mAbs, 05H02 and 2G7A1, approached the potency of potent spike targeting mAbs, and were >10-fold more potent than a previously reported anti-hACE2 antibody, h11b11, that has murine variable regions grafted onto a human antibody (Du et al., 2021).

**Table 1:** Antiviral potency (IC_50_ values, ng/ml) of fully human anti-hACE2 mAbs.

|  | **2C12H3** | **05G01** | **05H02** | **2G7A1** | **05B04** | **05B04**<br>**LC/**<br>**05D06**<br>**HC** |
| --- | --- | --- | --- | --- | --- | --- |
| **SARS-CoV-2 (Wuhan-Hu-1)** | 86.14 | 52.68 | 7.513 | 14.21 | 34.78 | 21.71 |
| **SARS-CoV-2 (Beta)** | 138.5 | 82.06 | 8.366 | 58.61 | 70.13 | 37.78 |
| **SARS-CoV-2 (Delta)** | 125.9 | 71.97 | 11.27 | 39.26 | 67.81 | 33.62 |
| **SARS-CoV-2 (Omicron)** | 98.13 | 74.17 | 8.49 | 17.97 | 56.65 | 22.13 |
| **SARS-CoV** | 76.57 | 34.67 | 7.563 | 9.602 | 31.54 | 16.26 |
| **Pangolin CoV-GD** | 102.9 | 31.28 | 6.329 | 6.595 | 37.03 | 20.35 |
| **Pangolin CoV-GX** | 82.89 | 66.09 | 7.823 | 18.23 | 41.01 | 28.06 |
| **Bat CoV Rs7327** | 29.47 | 21.85 | 14.37 | 3.848 | 12.32 | 13.47 |
| **Bat CoV Rs4231** | 44.14 | 33.02 | 2.221 | 4.139 | 37.6 | 13.38 |

IC<sub>50</sub> values for anti-hACE2 mAb inhibition of pseudotyped virus infection were determined using HIV-1 based pseudoviruses and Huh-7.5 target cells, as described in (Schmidt *et al.*, 2020), except that the anti-hACE2 mAbs were preincubated with cells rather than viruses prior to infection. 05B04 LC/ 05D06 HC is a hybrid mAb generated by coexpression of the 05B04 light chain with the heavy chain of a clonally related mAb, 05D06.

In addition to SARS-CoV-2 (Wuhan-hu-1), the anti-hACE2 mAbs inhibited infection by SARS-CoV-2 variant pseudotypes, including beta, delta, and omicron (BA.1), all with comparable potency (Figure 1B, Table 1). To further evaluate their antiviral breadth, anti-ACE2 mAb-treated Huh-7.5 cells were challenged with viruses pseudotyped with spike glycoproteins from SARS-CoV or SARS-related coronaviruses from other mammals, specifically Pangolin CoV-GD, Pangolin CoV-GX, Bat CoV Rs4231 and Bat CoV Rs7327. All of the sarbecoviruses tested were sensitive to inhibition by the human anti-hACE2 mAbs, with comparable potencies (IC_50_ values ranging from 2.2 ng/ml to 85.4 ng/ml) (Figure 1B, C, D, Table 1).

We next assessed the ability of four anti-human ACE2 mAbs to inhibit natural SARS-CoV-2 infection. Human Huh-7.5 or African green monkey (agm) Vero E6 cells were incubated with the the 05B04 mAb and challenged with SARS-CoV-2 USA_WA/2020. The 05B04 mAb inhbited infection with similar potency on both cell lines (Figure 2A). Because SARS-CoV-2 omicron/BA.1 replicates poorly in Huh-7.5 cells we next tested four antibodies using Vero E6 target cell only. Three mAbs, namely 2G7A1, 05B04 and 05B04LC/05D06HC potently inhibited both SARS-CoV-2 USA_WA/2020 and SARS-CoV-2 omicron/BA.1, while 05H02 was less potent in Vero E6 cells (Figure 2B) than predicted by its activity against SARS-CoV-2 spike pseudotypes in Huh-7.5 cells (Table 1). The agmACE2 protein shares 95% sequence similarity with hACE2, and sequence differences between hACE2 and agmACE2 might account for the reduced potency of 05H02 in Vero E6 cells (Figure 2B). Nevertheless, these findings demonstrate that human anti-hACE2 mAbs can inhibit infection by spike glycoprotein pseudotyped or authentic, replication-competent SARS-CoV-2.

**Figure 2.**
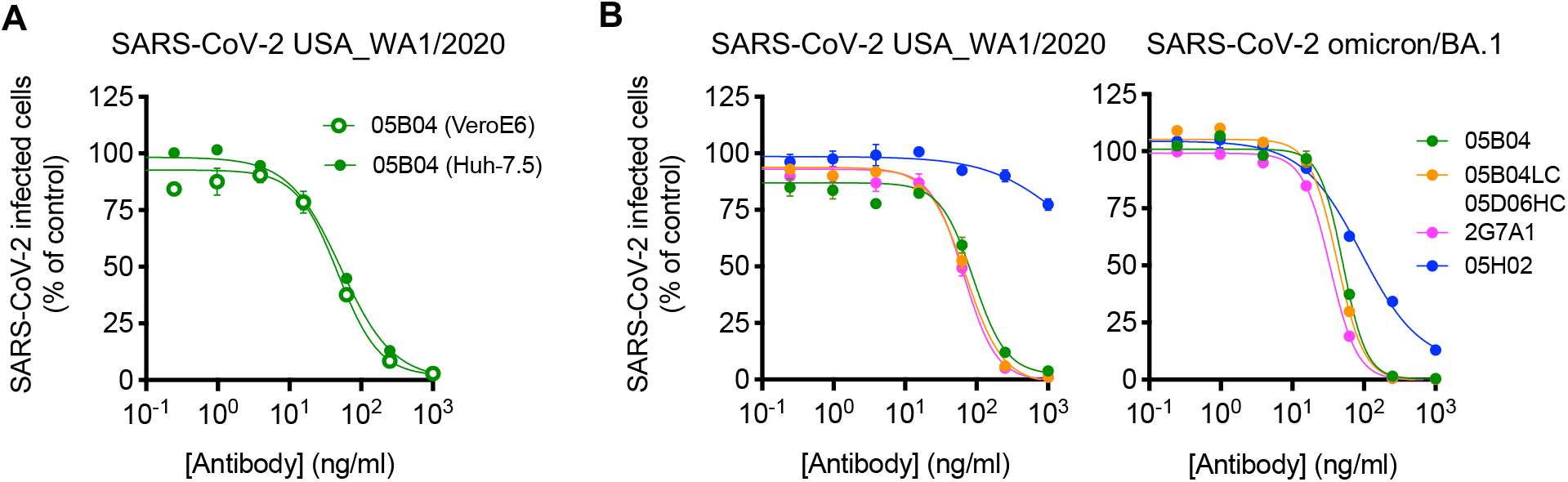
Inhibition of authentic SARS-CoV-2 infection by human anti-hACE2 mAbs. (A, B) Anti-hACE2 (hACE2) mAbs (2G7A1, 05H02, 05B04, and the hybrid antibody 05B04 LC/05D06 HC) were serially diluted and incubated with Vero E6 and Huh-7.5 target cells (A) or Vero E6 target cells only (B). Thereafter, cells were infected with authentic SARS-CoV-2 (WA1/2020) (A, B) or SARS-CoV-2 omicron BA.1 variants (B). Infected cells were quantified by immunostaining, and plotted as a percentage of the number of cells infected in the absence of anti-hACE2 mAbs. Mean and range of duplicate titrations.

### Human anti-hACE2 mAbs inhibit SARS-CoV-2 spike-hACE2 binding

We assessed the specificity, kinetics and affinity of the interaction between the human anti-hACE2 mAbs and hACE2, using flow cytometry and surface plasmon resonance (SPR) experiments. In flow cytometric analyses, each of the mAbs tested bound to A549 cells expressing human ACE2, but not to parental A549 cells (Figure 3A). Notably, each of the mAbs tested bound to A549 cells expressing macaque ACE2 and hACE2 equivalently (Figure S2). This property should facilitate the preclinical evaluation of the safety and efficacy of the mAbs in macaque models.

**Figure 3.**
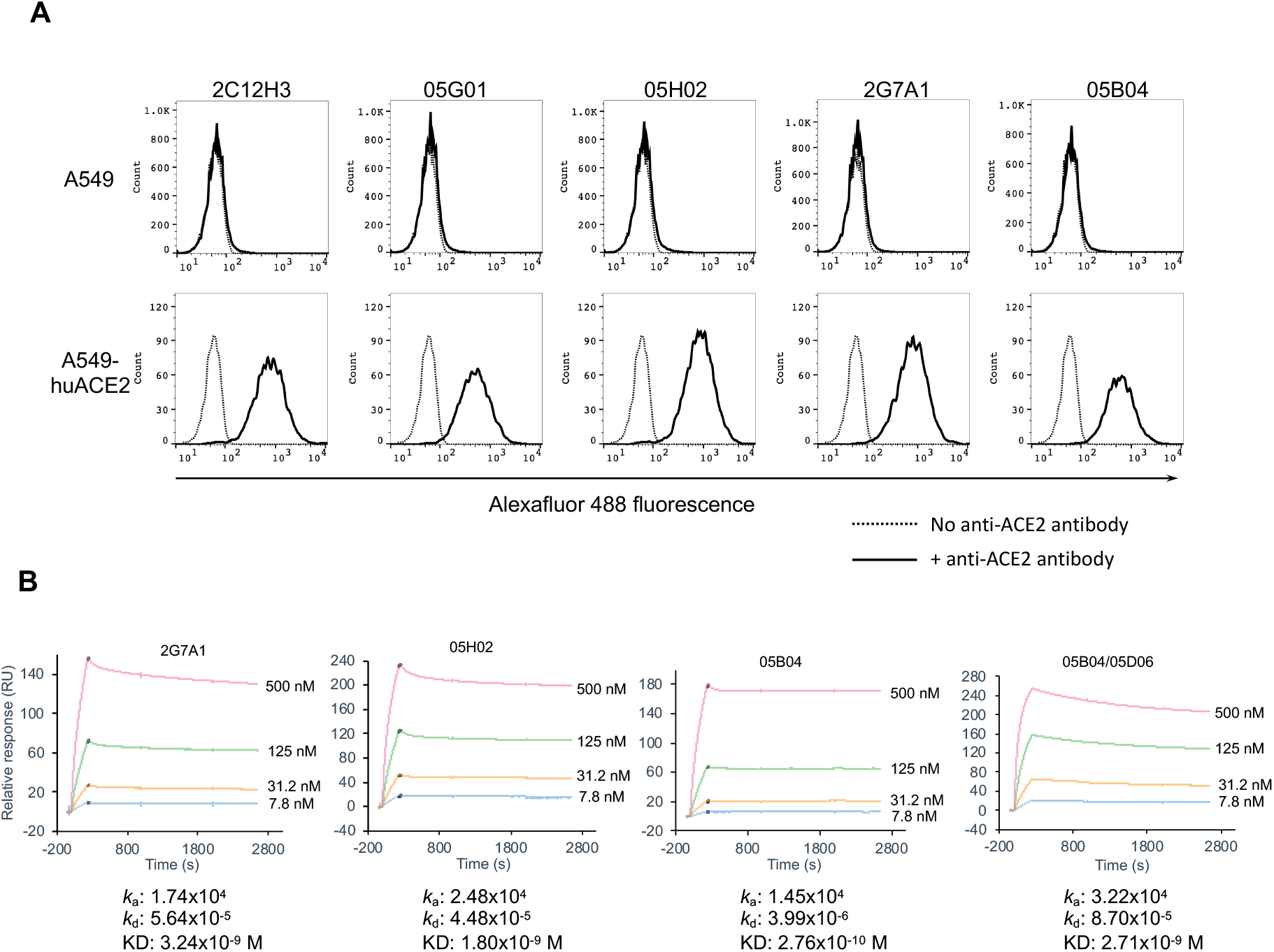
Binding to hACE2 by human anti-ACE2 antibodies. (A) Parental A549 cells (upper panels), or A549 cells stably expressing hACE2 (lower panels) were incubated in the presence (solid lines) or absence (dotted lines) of the indicated ACE2 antibodies. The cells were then incubated with Alexa Fluor 488 conjugated goat anti-human IgG and then analyzed by flow cytometry. (B) Four anti-hACE2 mAbs (2G7A1, 05H02, 05B04 and the hybrid antibody 05B04 LC/05D06 HC) were immobilized onto a Protein G Sensor chip. His-tagged ACE2 1-740 aa protein (7.8 nM, 31.2 nM, 125 nM, or 500 nM) were injected at 30 µl/min for 240 s followed by a dissociation phase of 2400 s at a flow rate of 30 µl/min. KD values were calculated from the ratio of association and dissociation constants (KD = *k*_d_/*k*_a_), derived using a 1:1 binding model.

In SPR experiments, four anti-hACE2 mAbs (2G7A1, 05H02, 05B04, and hybrid 05B04LC/05D06HC) were immobilized on the sensor chip, and then monomeric hACE2 was applied, thus the affinity measurements may underestimate the affinity of a bivalent mAb binding to a dimeric ACE2 on a cell surface. Nevertheless, the association constants between the mAbs and the soluble monomeric hACE2 ectodomain (1-740 aa) ranged from 1.45×10^4^ to 3.22×10^4^, while the dissociation constants ranged from 3.99×10^−6^ to 4.48×10^−5^ (Figure 3B). As such, the equilibrium dissociation constant (kD) values for the mAbs were in the nanomolar to picomolar range, specifically from 3.24×10^−9^ M to 2.76×10^−10^ M (Figure 3B).

We next developed a NanoLuc-fusion protein-based assay to measure recombinant hACE2 binding to trimeric SARS-CoV-2 spike proteins and its inhibition by the anti-hACE2 mAbs (Figure 4A). Specifically, we expressed prefusion conformationally stabilized secreted ‘HexaPro’ trimeric SARS-CoV2 spike proteins with NanoLuc luciferase fused at their C-termini (S-6P-NanoLuc). We then performed pulldown assays with His-tagged hACE2 immobilized on magnetic beads (Figure 4A). Preincubation of the immobilized hACE2 with four selected anti-hACE2 mAbs (2G7A1, 05H02, 05B04 and hybrid 05B04LC/05D06HC) blocked hACE2 binding to SARS-CoV-2 (Wuhan-Hu-1) trimers (Figure 4B). Moreover, and in agreement with results using the pseudotype neutralization assay (Figure 1B), the mAbs also inhibited omicron BA.1 spike trimer binding to immobilized hACE2 with similar potency (Figure 4B). Overall, we conclude that human anti-ACE2 mAbs block binding interactions between hACE2 and SARS-CoV-2 spike and consequently inhibit spike glycoprotein virus infection with breadth and, in particular, in the case of 2G7A1, 05H02, 05B04, and 05B04LC/05D06HC, high potency.

**Figure 4.**
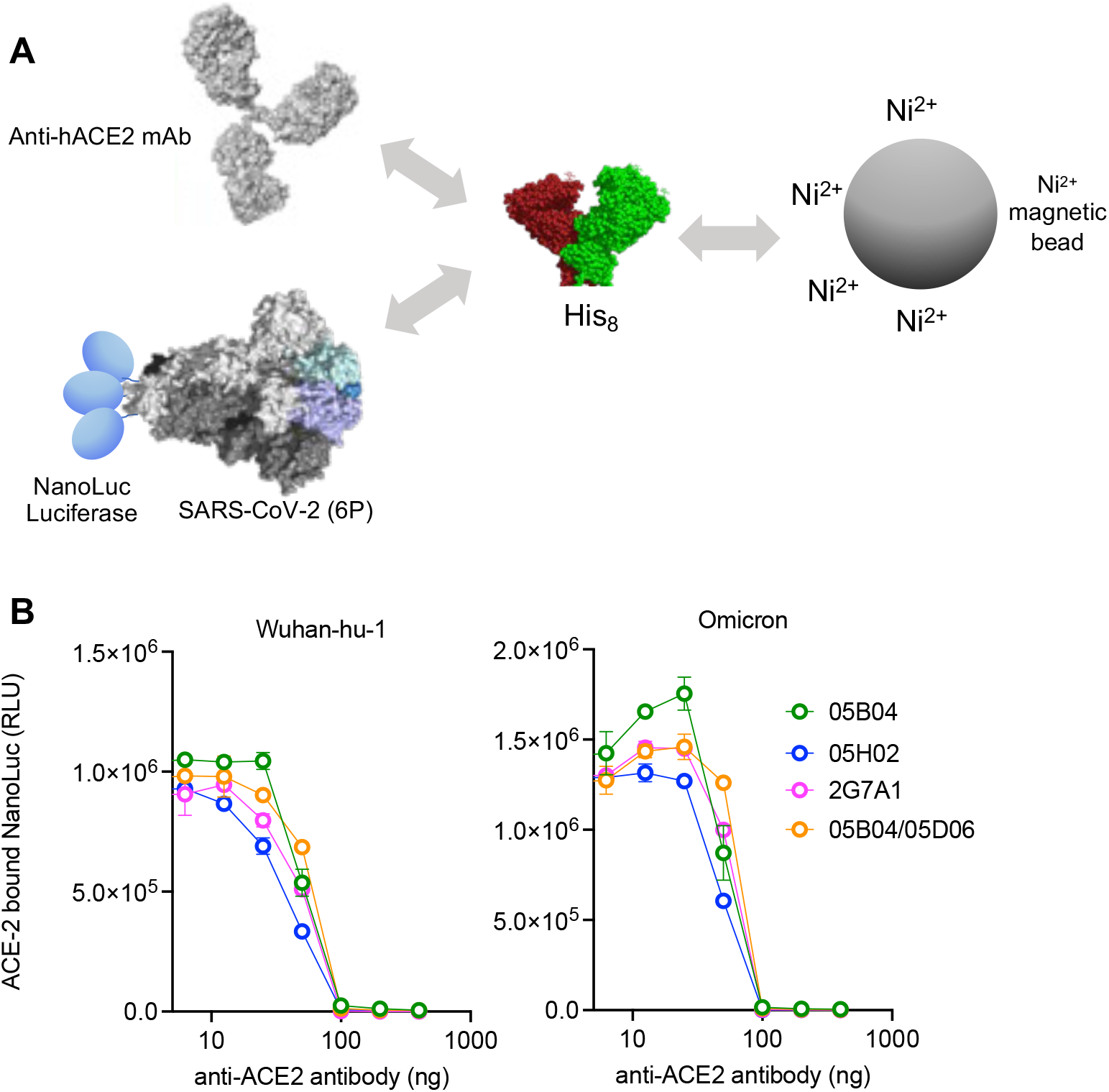
Inhibition of ACE2:Spike binding by anti-hACE2 antibodies. (A) Schematic representation of the spike-hACE2 binding assay in which NanoLuc luciferase was appended to the C-termini of a conformationally stabilized SARS-CoV-2 spike trimer, S-6P-NanoLuc (based on Wuhan-hu-1, or omicron BA.1 variants). The fusion protein was incubated with His-tagged ACE2 (1-740 aa) that was preincubated in the presence or absence of anti-hACE2 mAbs, and complexes were captured using His-tag magnetic beads. (B) Serially diluted mAbs (2G7A1, 05B04, 05H02, or 05B04LC/05D06HC) were mixed with 100 ng of His-tagged ACE2 1-740 aa proteins. After incubation, the mixture was then incubated with Wuhan-hu-1 or Omicron S-6P-NanoLuc proteins (10 ng) followed by capture on His-tag magnetic beads. Bound NanoLuc activity was measured after washing.

### Structural analysis of anti-hACE2 mAbs

To delineate the structural basis for broad neutralization of anti-hACE2 mAbs, we determined the structure of soluble hACE2 (Chan et al., 2020) bound to the 05B04 Fab fragment using single-particle cryo-electron microscopy (cryo-EM). Focused refinement resulted in a 3.3Å resolution map, revealing a 05B04 Fab bound to the N-terminal helices of hACE2 (Figure 5A,B, Figure S3A-D and Table S3). The binding orientation of 05B04 sterically-hinders and competes with SARS-CoV-2 RBD (Figure 5C) and is similar to the binding orientations and epitopes of murine antibodies 3E8 and h11b11 (Figure S3E-G) (Chen *et al*., 2021; Du *et al*., 2021). The 05B04 complementary determining region (CDR) loops CDRH2, CDRH3, CDRL1, and CDRL3 contribute to binding an epitope that comprises residues from the α1 and α2 helices, resulting in a total buried surface area (BSA) of ∼1280 Å^2^ (620 Å^2^ epitope BSA + 660 Å^2^ paratope BSA). The CDRH3 loop mediates extensive polar and van der Waals contacts, including insertion of Met98_HC_ into a hydrophobic pocket involving Phe28, Leu79, Met82, and Tyr83 of ACE2 (Figure 5D). CDRH2, CDRL1, and CDRL3 loops contribute to a hydrogen bond network with both main chain and side chain atoms of ACE2. For example, Gln24 of ACE2 forms a hydrogen bond with the hydroxyl group of Tyr32_LC_, while ACE2 residue Glu23 forms a salt bridge with Arg56_HC_ (Figure 5E). These interactions mimic favorable interactions between the SARS-CoV-2 RBD ridge and the N-terminal α1 helix of ACE2, which were critical to increases in affinity for SARS-CoV-2 RBD relative to SARS-CoV RBD (Shang et al., 2020; Yan et al., 2020). Thus, 05B04-mediated inhibition of ACE2-binding sarbecoviruses is achieved through molecular mimicry of select SARS-CoV-2 RBD interactions, providing high binding affinity to ACE2 despite the smaller binding footprint on ACE2 relative to the RBD (Figure S3F,G).

**Figure 5.**
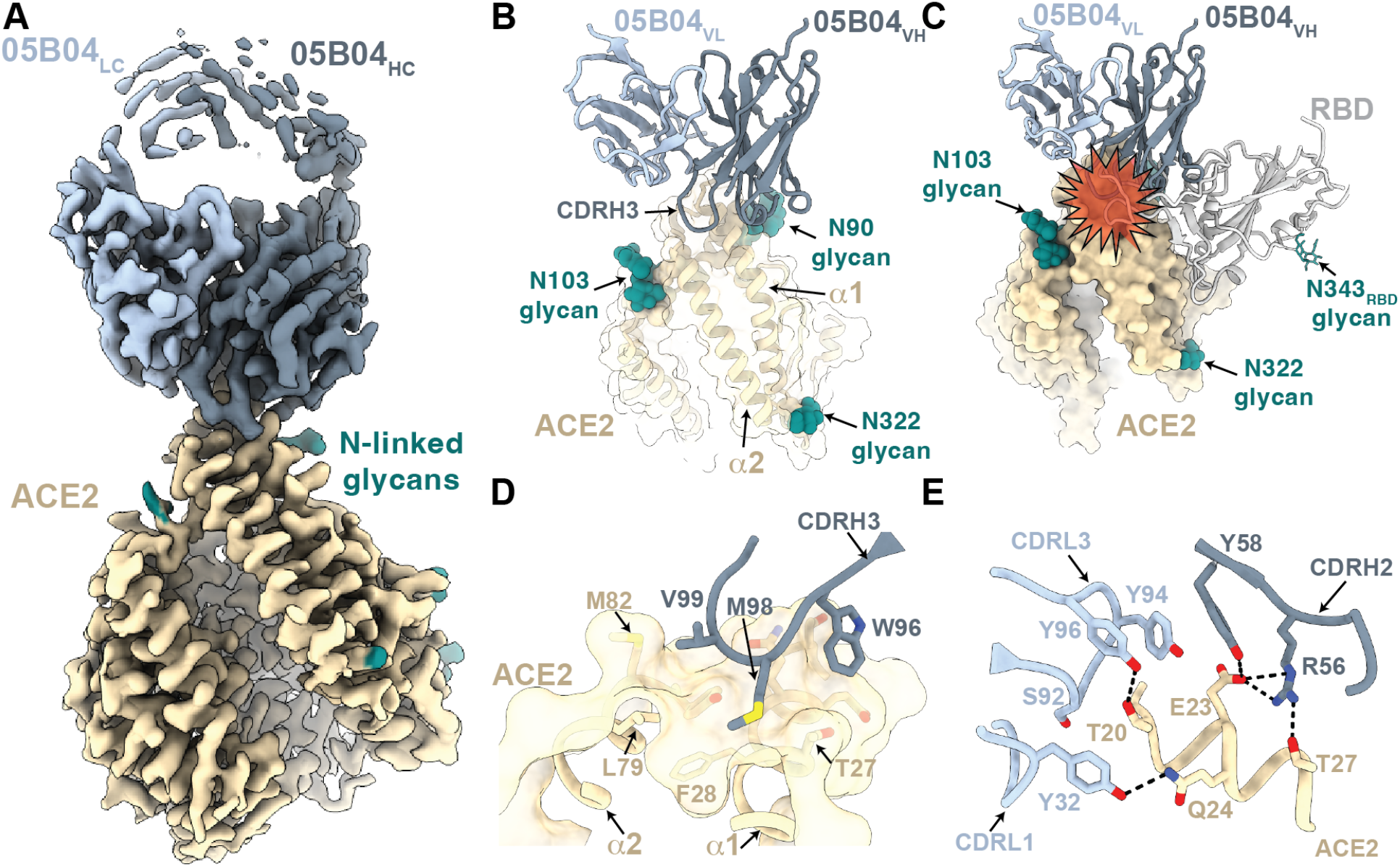
Cryo-EM structure of the 05B04-hACE2 complex. (A) 3.3Å cryo-EM density for the 05B04 Fab – hACE2 complex. Density for 05B04 Fab variable domains are shown as shades of slate blue and hACE2 is shown as wheat. (B) Close-up view of 05B04 variable domains (blue cartoon) binding to hACE2 (surface and cartoon, wheat). (C) Superimposition of PDB 6VW1 on the 05B04-ACE2 structure, aligned on the Cα atoms of the ACE2 α1 and α2 helices. Predicted clashes between the RBD (gray) and 05B04 variable domains (shades of blue) are highlighted with a red star. (D) CDRH3-mediated contacts on ACE2 (surface and cartoon rendering). (E) CDRH2, CDRL1, and CDRL3-mediated contacts with the α1 helix. Potential hydrogen bonds are shown as black dashed lines.

### Anti-hACE2 mAbs do not inhibit hACE2 activity or induce hACE2 downregulation

ACE2 is a zinc metalloprotease whose physiological function is to catalyze the hydrolysis of angiotensin I to angiotensin 1-9, or angiotensin II to angiotensin 1-7 (Donoghue et al., 2000; Tipnis et al., 2000). The ACE2 enzyme active site (Towler et al., 2004) is distal to the N-terminal α1 helix, where residues that contact sarbecovirus spike proteins are primarily located (Lan et al., 2020) and 05B04-like antibodies bind (Figure S3E-G). Nevetheless, it was possible that binding of the anti-ACE2 antibodies to ACE2 could allosterically hinder ACE2 enzymatic activity. To measure the impact of the anti-hACE2 mAbs on ACE2 activity, anti-ACE2 mAbs 2G7A1, 05B04, 05H02, and hybrid mAb 05B04LC/05D06HC were mixed with ACE2 protein at various molar ratios (10:1, 50:1, or 250:1) and ACE2 enzymatic activity determined. While the ACE2-specific inhibitor MLN-4760 inhibited ACE2 activity at concentrations of 10-100 nM (Figure 6A), none of anti-ACE2 mAbs tested (2G7A1, 05B04, 05H02, and 05B04LC/05D06HC) affected ACE2 activity, even when present at concentrations representing a 250-fold molar excess over the hACE2 protein (Figure 6B). Thus, while allosteric inhibition of ACE2 activity by the mAbs was theoretically possible, these results suggest that such inhibition does not occur.

**Figure 6.**
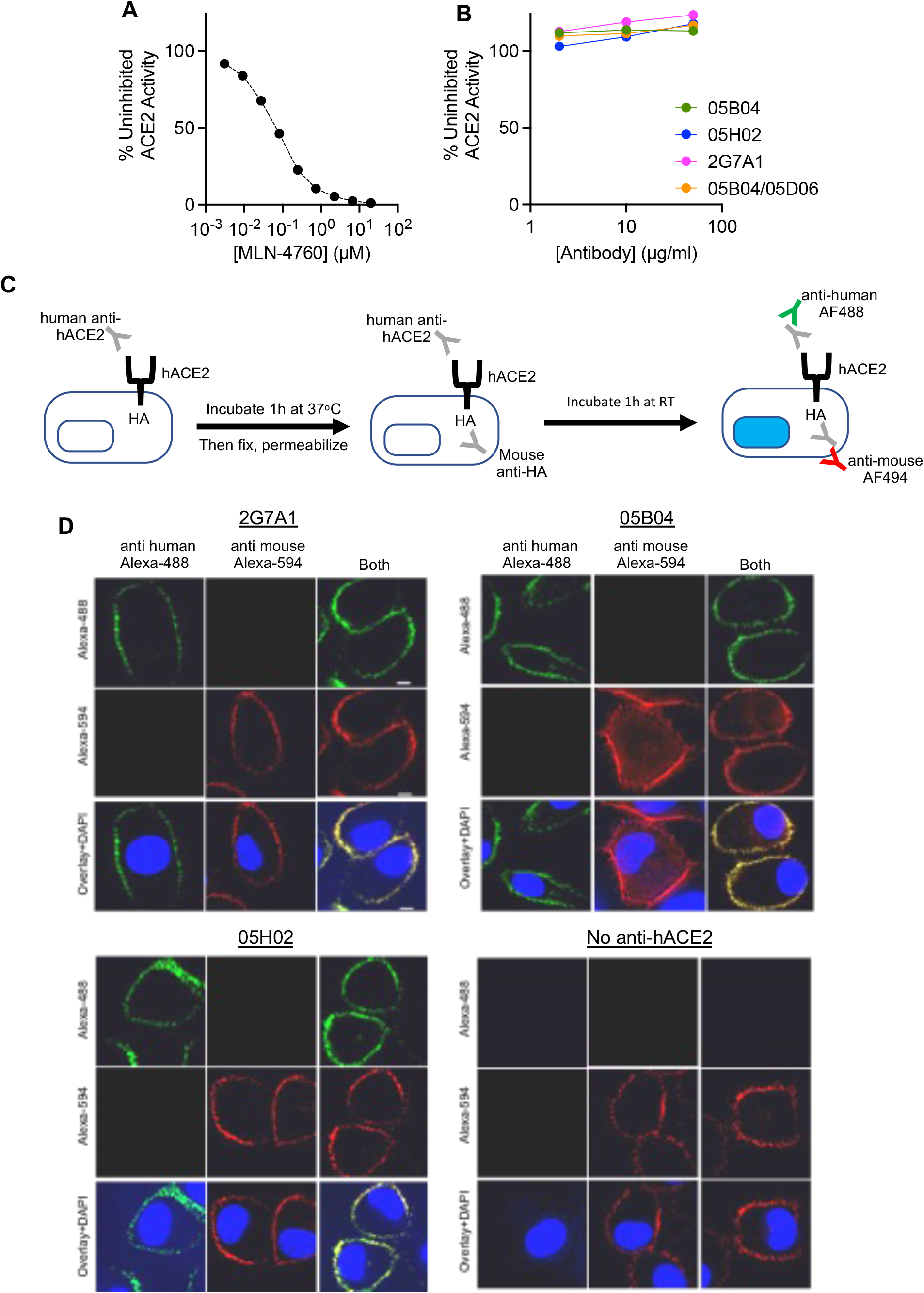
No effect of anti-hACE2 mAbs on hACE2 activity and distribution. (A, B) the ACE2 inhibitor control (MLN-4670, A) or anti-hACE2 mAbs (2G7A1, 05B04, 05H02, and 05B04LC/05D06HC, 2 μg/ml, 10 μg/ml, 50 μg/ml (B) were mixed with ACE2 (0.2 μg/ml) and a fluorogenic ACE2 substrate in 96-well plates. After incubation, fluorescence intensity (555 nm/585 nm, excitation/emission), indicative of ACE2 enzymatic activity, was measured and plotted as a percentage of the uninhibited control. (C) Outline of an assay to determine hACE2 and anti-hACE2 mAb internalization. Live A549 cells expressing a C-terminally (intracellular) HA-tagged hACE2 receptor were incubated with anti-hACE2 mAbs, Then, cells were fixed and permeabilized. Total hACE2-HA was then immunostained with a mouse anti-HA-tag antibody. The internalization of hACE2-HA and anti-hACE2 mAbs was then evaluated by staining with anti-mouse Alexa Fluor 594 and goat anti-human Alexa Fluor 488, respectively. (D) Localization of anti-hACE2 mAbs (green, left) or hACE2-HA (red, center) or both (right) following incubation of live A549/ACE2-HA cells with 2G7A1, 05B04, or 05H02, or the absence of human anti-ACE2 antibody as indicated. Blue stain (DAPI) indicates cell nuclei.

ACE2 is an integral membrane protein, with the catalytic domain anchored to the exterior of cells via C-terminal transmembrane and intracellular domains. Antibody binding to membrane proteins can induce their internalization and thereby reduce cell surface levels. To determine whether anti-hACE2 mAb binding induced hACE2 internalization, we developed a fluorescence-based assay in which anti-hACE2 mAbs were incubated with A549 cells expressing hACE2 that was HA-epitope tagged at its intracellular C-terminus. After incubating the cells with anti-hACE2 mAbs for 1 hr at 37°C, cells were fixed and then permeabilized. Thereafter, the anti-hACE2 mAbs and the HA-tagged hACE2 were stained to reveal their subcellular distribution (Figure 6C). As a control, cells were incubated with a labelled anti-CD44 antibody, which is known to be internalized in A549 cells (Glatt et al., 2016). Each of anti-ACE2 mAbs tested (2G7A1, 05B04, and 05H02) exhibited near complete colocalization with hACE2-HA marked by anti-HA antibody staining. Importantly, both the hACE2 protein, as detected by anti-HA antibody, and the anti-hACE2 mAbs, detected with anti-human IgG antibody, remained localized on the cell surface, while the control anti-CD44 antibody accumulated at intracellular sites (Figure 6D, Figure S4). Indeed, only very low levels of intracellular hACE2 were detected, irrespective of mAb treatment, indicating that the anti-hACE2 mAbs had no effect on ACE2 internalization or recycling. This finding also suggested that the anti-hACE2 mAbs would be unlikely to undergo accelerated target-dependent clearance from the circulation during *in vivo* use.

### hACE2 mAbs protect hACE2 knock-in mice against SARS-CoV-2 infection

We next determined whether the human anti-hACE2 mAbs could protect against SARS-CoV-2 infection in an animal model. We used hACE2 knock-in mice for these experiments, in which the endogenous mouse ACE2 is replaced by hACE2, as the hACE2 is more likely to mimic the levels and distribution of ACE2 expression encountered in humans, compared to hACE2 transgenic mice. Importantly, hACE2 knock in mice are susceptible to SARS-CoV-2 infection (Zhou et al., 2021).

First, we determined the pharmacokinetic behavior of the hACE2 mAbs in the hACE2 knock-in mice. To measure the levels of hACE2 mAb in mouse serum, we generated a hACE2 (1-740)-NanoLuc fusion protein that could bind to anti-hACE2 mAbs and then be captured by protein G magnetic beads (Figure S5A). This assay allowed the detection of ∼0.01 ng to 10 ng of anti-ACE2 mAb in mouse serum, with linear standard curves in this concentration range (Figure S5B). Mice were injected subcutaneously with 250 μg of each anti-hACE2 mAb (∼12.5 mg/kg) on day 0 and the levels of mAbs in serum were measured until day 14. For 3 out of the 4 mAbs, namely 05B04, 05H02 and 2G7A1, the serum mAb concentration remained above 10 μg/ml on day 14 (Figure 7A), a value that is 100 to 1000-fold higher than their IC_50_ values. For 05B04LC/05D06HC, the mAb levels remained stable for around 7 days but dropped more dramatically from day 7 to 14, suggestive of the onset of an immune response to the non-self human antibody. The mean serum half-life for 05B04, 05H02 and 2G7A1 over 14 days was 8.3d, 5.3d, and 9.6d, which is similar to that typically observed for human IgG1 in mice (Vieira and Rajewsky, 1988). Overall, the anti-hACE2 mAbs showed favorable pharmacokinetics and conferred no obvious ill effects on the hACE2 knock-in mice.

**Figure 7.**
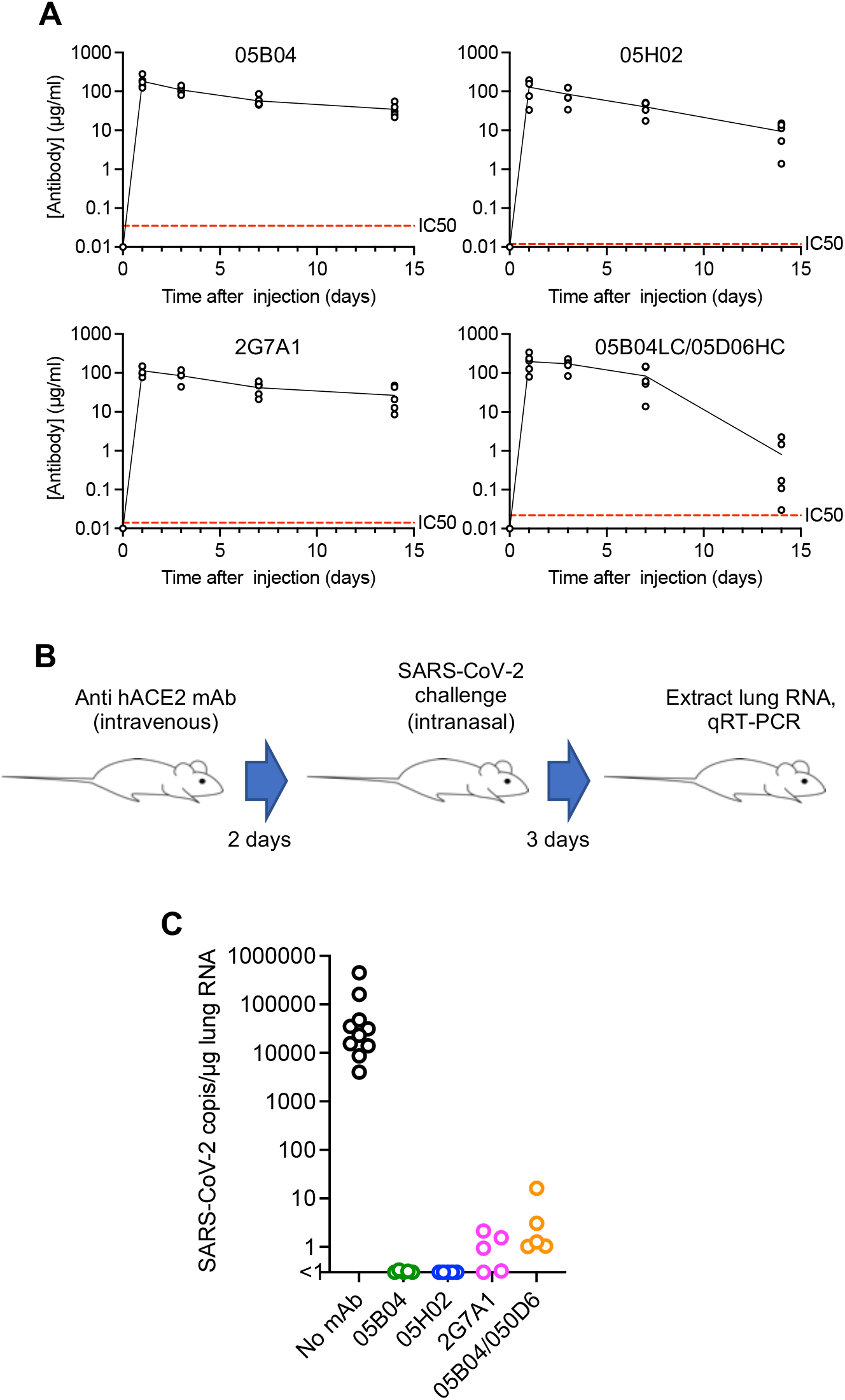
Evaluation of human anti-ACE2 antibodies as prophylaxis agents in mice. (A) Anti-hACE2 mAb levels following subcutaneous injection of 250 μg (equivalent to approximately 12.5 mg/kg) of anti-hACE2 antibodies (05B04, 05H02, 2G7A1, and 05B04LC/05D06HC) into each of five hACE2 knock-in mice (B6.129S2(Cg)-Ace2tm1(ACE2)Dwnt/J,) on day 0. Serum dilutions (0.5 μl, 0.1 μl, or 0.02 μl) of the sera collected on the indicated days or mAb standards were mixed with ACE2 (1-740 aa)-NanoLuc fusion protein and captured by Protein G beads (see Figure S4). Dashed red line = IC_50_ for inhibition of SARS-CoV-2 pseudotyped virus infection. (B, C) Human ACE2 knock-in mice (B6.129S2(Cg)-Ace2tm1(ACE2)Dwnt/J,) (n=5) were injected with 250 μg (equivalent to 12.5 mg/kg) of the anti-human ACE2 antibodies (05B04, 05H02, 2G7A1, or 05B04LC/05D06HC). At 2 days after antibody injection mice were challenged intranasally with SARS-CoV-2, USA_WA/2020 P3, 2×10^5^ PFU/mouse (virus titers measured on VeroE6 cells) (B). Three days later after mouse lungs were harvested and RNA extracted. The number of viral genomes per microgram of total lung RNA was measured using qRT-PCR (C).

We next asked whether the anti-ACE2 mAbs could provide protection against SARS-CoV-2 infection. Each of the four anti-hACE2 mAbs were injected intravenously into mice (5 mice per group) two days prior to intranasal challenge with 2×10^5^ PFU of human SARS-CoV-2, strain USA_WA/2020 P3 (Figure 7B). Levels of viral RNA in the lungs of mock treated mice were in the range of 10^4^ to 10^6^ copies/μg total RNA at 2 days after infection, comparable to previous observations using this model (Zhou *et al*., 2021). Strikingly, pre-treatment of the mice with any one of the four anti-hACE2 mAbs, 2G7A1, 05B04, 05H02, or 05B04LC/05D06HC dramatically reduced SARS-CoV2 replication in lungs, to levels below or close to the limit of detection (∼1 copy of viral RNA/μg total RNA) (Figure 7C). Therefore, when used prophylactically the anti-hACE2 mAbs provided near sterilizing protection against lung SARS-CoV-2 infection in hACE2 knock-in mice.

## Discussion

As SARS-CoV-2 has adapted to the immunological pressures imposed by naturally elicited spike-binding antibodies, therapeutic mAbs derived from natural human anti-spike repertoires have lost effectiveness and are now largely obsolete (Cao *et al*., 2022). While it would likely be possible to isolate second, and perhaps multiple generations of spike-targeting human mAbs, the threat of viral resistance will remain, as SARS-CoV-2 continues to adapt to the range of human antibodies it encounters in previously infected or vaccinated individuals. Thus, the current pace of SARS-CoV2 evolution makes the continued derivation of anti-spike therapeutic mAbs an impractical way of combating SARS-CoV-2. Human anti-hACE2 mAbs offer the possibility of durable, ‘resistance proof’ prophylaxis and treatment. While it is conceivable that SARS-CoV-2 might acquire the ability to use alternative receptors (Puray-Chavez et al., 2021), such an event would represent a much greater genetic hurdle than does the evasion of spike-targeting mAbs.

ACE2 plays a role in the regulation of cardiovasular and renal function (Donoghue *et al*., 2000; Hamming et al., 2007), and mAbs targeting ACE2 have the potential to exhibit unwanted side effects. Notably however, the ACE2 enzymatic active site is distal to the site that is bound by sarbecovirus spike proteins (Lan *et al*., 2020; Towler *et al*., 2004) and the anti-hACE2 mAbs described herein. Structural comparisons between mAb 05B04 and the SARS-CoV-2 RBD reveal epitope competition and steric hinderance as the antiviral mechanism, and no antibody-induced allosteric changes in the enzyme active site. Indeed, none of the mAbs described herein affected ACE2 enzymatic activity and, moreover, none induced the internalization of ACE2 that is normally localized on the cell surface. Thus, we do not expect the mAbs to have deleterious side effects, based on their target specificity.

Anti-hACE2 mAbs should be of particular utility in human populations with immunodeficiency or undergoing immunosuppressive treatment who are especially vulnerable to severe outcomes from SARS-CoV-2 infection and in which vaccine-elicited protective immunity is more difficult to achieve (Akalin et al., 2020; Deepak et al., 2021; Prendecki et al., 2021). MAbs with the half-life extending ‘LS’ mutations are expected to provide prophylactic protection for weeks or months (Zalevsky *et al*., 2010), and would likely enable a single protective administration during the early phase of the intermittent waves of high incidence that have thus far characterized the SARS-CoV-2 pandemic. Moreover, the exceptional breadth and potency with which these mAbs are able to inhibit infection by ACE-2 binding sarbecovirus infection should make them uniquely effective interventions in the event that future outbreaks result from spillover of this group of pandemic threat coronaviruses.

## Acknowledgements

The sACE2 WT plasmids used in the CyoEM experiments were a gift from Erik Procko (Addgene plasmid # 145147). Cryo-EM data for this work was collected at the Stanford-SLAC cryo-EM center with support from Dr. Elizabeth Montabana. This work is supported by grants from NIAID: P01 AI165075 (P.D.B., T.H. and C.M.R.) R37AI64003 to P.D.B., R01AI78788 to T.H.. C.O.B. is supported by the Howard Hughes Medical Institute Hanna Gray Fellowship and is a Chan Zuckerberg Biohub investigator. PDB is a Howard Hughes Medical Institute Investigator. This article is subject to HHMI’s Open Access to Publications policy. HHMI lab heads have previously granted a nonexclusive CC BY 4.0 license to the public and a sublicensable license to HHMI in their research articles. Pursuant to those licenses, the author-accepted manuscript of this article can be made freely available under a CC BY 4.0 license immediately upon publication.

## Author Contributions

F.Z. T.H. and P.D.B. conceived the study. F.Z and J.J. constructed, expressed and purified immunogens and human anti ACE2 antibodies and performed the pseudovirus antiviral assays, affinity measurements, ACE2 competition assays, flow cytometry assays, ACE2 activity and internalization assays and the pharmacology experiments. R.V.H.dC. S. N-H., and F.Z. did in vivo prophylaxis experiments. E.N., D.A., I.L., I.C.L., did the AlivaMab immunizations, hybridoma isolation and ACE2 binding screen. T.H. and H-H.H. did the in vitro live virus inhibition assays. T.C., M.E.A., and C.O.B., designed and carried out cryo-EM experiments. P.D.B., T.H., E.N., C.M.R., G.V. and C.O.B. supervised the work. F.Z., C.O.B., T.H. and P.D.B. wrote the manuscript with input from all co-authors.

## Data Availability

The atomic model and cryo-EM map generated for the 05B04-hACE2 complex has been deposited at the Protein Databank (PDB) (http://www.rcsb.org/) and the Electron Microscopy Databank (EMDB) (http://www.emdataresource.org/) under accession codes 8E7M and EMD-27939, respectively.

## Declaration of Interests

The Rockefeller University has filed a patent application for anti-hACE2 antibodies on which F.Z., T.H. and P.D.B. are listed as inventors.

## Methods

### Cell lines

Human embryonic kidney HEK-293T cells (ATCC CRL-3216) and the derivative expressing ACE2, ie 293T/ACE2.cl22 (Schmidt *et al*., 2020), Caco-2 cells (ATCC HTB-37^™^), human hepatoma-derived Huh-7.5 cells (Blight et al., 2002), Vero E6 cells and a derivative expressing TMPRSS2 (Wang et al., 2022) and A549 cells (adenocarcinomic human alveolar basal epithelial cells) were maintained in Dulbecco’s Modified Eagle Medium (DMEM) supplemented with 10% fetal bovine serum (Sigma F8067) and gentamycin (Gibco). All cell lines used in this study were monitored periodically to ensure the absence of retroviral contamination and mycoplasma.

### Generation of hACE2 specific human mAbs

ACE2-binding mAbs with human variable regions were generated using AlivaMab Mouse (Ablexis LLC) transgenic mouse strains. Specifically, AlivaMab mice were immunized subcutaneously with recombinant human ACE2 extracellular domain (1-740aa) fused to human IgG1 Fc and/or a poly-histidine tag. Mice were immunized at 3-week intervals for at least 4 times, using 10 µg subcutaneous injections at different sites. Mice with sera exhibiting SARS-CoV-2 infection inhibiting activity in the pseudotype virus assay were acute boosted prior to fusion of splenocytes with SP2/0 cells for hybridoma generation. Hybridomas expressing antibodies that bound to ACE2 were identified by enzyme-linked immunosorbent assay (ELISA) using plates coated with purified human ACE2. Anti-ACE2 hybridoma supernatants that contained antibodies with human ACE2 binding activity were tested for inhibition of SARS-CoV-2 pseudotyped virus infection of Huh-7.5 cells. This analysis indicated ten hybridoma antibodies that were positive for binding and potently inhibited SARS-CoV-2 pseudotype virus infection that were chosen for hybridoma cell subcloning and expansion. Antibodies were purified from these hybridoma culture supernatants and were further tested for potency ranking in the SARS-CoV-2 pseudotype virus inhibition assay.

### Human mAb expression plasmids

DNA encoding the variable regions of the heavy (VH) and light (VL) from hybridomas was PCR amplified from DNA extracted from the hybridoma cell lines. For the 05B04 and 05D06 antibodies, the DNA sequences encoding VH and VL were human codon-optimized using GenSmart™ Codon Optimization, synthesized by IDT. For each mAb, DNA encoding VH were fused to cDNA encoding the Fc domain of human IgG1, in which Fc domain was modified to include the substitutions at L234 L235 (LALA) that abolish FcR-gamma interaction, and substitutions at M428 N434 (LS) that enhance interaction with the neonatal Fc receptor to prolong mAb half-life in humans (Tamm and Schmidt, 1997; Wang *et al*., 2018; Zalevsky *et al*., 2010). To construct the expression plasmids for heavy and light chain antibody expression (von Boehmer et al., 2016), PCR amplicons or synthetic DNA encoding variable regions were subcloned using AgeI and XhoI (for LC), or AgeI and SalI (for HC), respectively, using NEBuilder® HiFi DNA Assembly.

### Human ACE2 and sarbecovirus spike expression plasmids

Plasmids expressing the spike proteins from SARS-CoV, SARS-CoV-2 (Wuhan-hu-1, Beta (B.1.351), Delta (B.1.617.2), and Omicron (B.1.1.529) variants), the pangolin (*Manis javanica*) coronaviruses from Guangdong, China (pCoV-GD) and Guanxi, China (pCoV-GX) were previously described (Schmidt *et al*., 2021a; Schmidt *et al*., 2020; Schmidt et al., 2021b). Human codon-optimized cDNAs encoding spike proteins from the rufous horseshoe bat (*Rhinolophus sinicus*) coronaviruses Rs4231 and Rs7327 were generated using GenSmart™ Codon Optimization, synthesized by IDT as gBlocks, and inserted into the pCR3.1 expression vector using NheI and XbaI and NEBuilder® HiFi DNA Assembly.

To construct the plasmids expressing NanoLuc-fused to conformationally stabilized versions of the SARS-CoV-2 Wuhan-hu-1 or Omicron variant spike proteins, the HexaPro (6P) modified cDNAs was fused at its C-terminus with DNA encoding a trimerization domain, a GGSGG spacer sequence, NanoLuc luciferase (NLuc), a human rhinovirus 3C protease cleavage site and a polyhistidine tag (8XHis). This cDNA, termed (S-6P-NanoLuc) was inserted into the pCR3.1 expression vector.

To construct a plasmid expressing catalytically inactive, His-tagged, or IgG1 Fc fused soluble ectodomain of ACE2 (1-740 aa), H374N and H378N substitutions were introduced by overlap extension PCR into a hACE2 cDNA and, an His-tag was fused to its C-terminus, and the purified PCR product was inserted into the pCAGGS expression vector (ACE2-1-740aa-His). To construct the expression plasmid encoding hACE2 (1-740aa)-NanoLuc-8XHis, the ACE2 1-740aa and NanoLuc-8XHis fragments were PCR amplified using ACE2-1-740aa-His and S-6P-NanoLuc as templates, respectively, followed by Gibson assembly and insertion into the pCR3.1 expression vector.

### Protein and antibody expression and purification

To express the momomeric, His-tagged human ACE2 extracellular domain (residues 1-740) used as immunogen or soluble hACE2 used for cryo-EM studies (residues 1-614), Expi293 cells were transfected with the expression plasmid hACE2-1-740aa-8xHis or hACE2-1-614aa-8xHis using ExpiFectamine 293 (ThermoFisher Scientific). Four days later, the supernatant was filtered with 0.22-um membrane filter and loaded on Ni-NTA agarose (Qiagen) and, after washing, ACE2 proteins were eluted with 200 mM imidazole in PBS. For cryo-EM structural studies, a subsequent size-exclusion chromatography step on a superdex200 10/300 column (Cytiva) was performed against PBS, and fractions corresponding to monomeric soluble ACE2 were pooled and stored at 4°C. Dimeric, Fc-fused ACE2 extracellular domain was also expressed in Expi293 cells in the same way. The secreted proteins in supernatant were first incubated with Protein G Sepharose 4 Fast Flow overnight at 4°C, loaded into column, and, after washing, eluted with 0.1 M glycine, pH 2.9 into tubes containing 1/10th volume of 1 M Tris, pH8.0.

To express mAbs, Expi293 cells were transfected with the corresponding light chain and heavy chain expression plasmids at the ratio of 1:1 using ExpiFectamine 293. Four days later, the mAbs in the supernatant were purified through Protein G Sepharose 4 Fast Flow and eluted with 0.1 M glycine, pH 2.9 as described above.

To express S-6P-NanoLuc proteins, Expi293 cells were transfected with S-6P-NanoLuc expression plasmids that included for the original Wuhan-hu-1 or Omicron spike variants using ExpiFectamine 293. Three days later, the supernatant was harvested and loaded on Ni-NTA agarose and, after thorough wash, S-6P-NanoLuc proteins were released after HRV 3C protease (TaKaRa) treatment overnight at 4°C.

To express the His-tagged ACE2 (1-740aa)-NanoLuc proteins, Expi293 cells were transfected with expression plasmids ACE2(1-740aa)-NanoLuc-8XHis using ExpiFectamine 293. Four days later, the supernatant was harvested and loaded on Ni-NTA agarose and, after washing, ACE2(1-740aa)-NanoLuc proteins were eluted with 200 mM imidazole in PBS. All recombinant proteins, including purified mAbs, were dialyzed against PBS before used in further experiments.

### Sarbecovirus spike-bearing pseudotypes and infectivity inhibition assay

To generate HIV-1 virions pseudotyped with Sarbecovirus spikes, including SARS-CoV, SARS-CoV-2 (Wuhan-hu-1, Beta, Delta, and Omicron variants), pangolin coronavirus pCoV-GD, pangolin coronavirus pCoV-GX, bat coronaviruses:, bat SARSr-CoV 4231, and bat SARSr-CoV 7327, ten million 293T cells in a 15 cm dish were transfected with 25 μg of an HIV-1 envelope-deficient proviral plasmid expressing NanoLuc along with 7.5 μg of spike expression plasmids, in which the C terminal 19aa was truncated (Δ19) (Schmidt *et al*., 2020). Cells were washed twice with PBS the next morning and virions were harvested at 48 hr post transfection, filtered (0.22 μm), and purified by Lenti-X Concentrator (TaKaRa). To measure the infectivity, viral stocks were two-fold serially diluted and added to Huh-7.5 cells, a SARS-CoV-2 susceptible cell line, in 96-well plates seeded one day prior to infection. Cells were then harvested at 48 hr post infection for measuring NanoLuc activity using the Nano-Glo Luciferase Assay System and GloMax® Navigator Microplate Luminometer (Promega).

To measure antiviral activity, the hACE2 mAbs were four-fold serially diluted (beginning with 2 μg/ml) in 96-well plates over seven dilutions and incubated with Huh-7.5 target cells for 1 hr at 37°C. Thereafter, the mAb-treated Huh-7.5 cells were infected with sarbecovirus spike pseudotyped viruses. Cells were harvested 48 hr post infection and NanoLuc luciferase activity measured in infected cells as described above.

### SARS-CoV-2 virus stocks and titration

SARS-CoV-2 strains USA-WA1/2020 and the Omicron variant B.1.1.529 were obtained from BEI Resources (catalog no. NR-52281 and NR-56461, respectively). The original WT virus (WA1/2020) was amplified in Caco-2 cells, which were infected at a multiplicity of infection (MOI) of 0.05 plaque forming units (PFU)/cell and incubated for 5 days at 37^ο^C. The Omicron variant B.1.1.529 was amplified in Vero E6 cells obtained from the ATCC that were engineered to stably express TMPRSS2. Vero-TMPRSS2 cells were infected at a MOI = 0.05 PFU/cell and incubated for 4 days at 33°C. Virus-containing supernatants were subsequently harvested, clarified by centrifugation (3,000 *g* × 10 min), filtered using a disposable vacuum filter system with a 0.22 μm membrane and stored at −80°C. Virus stock titers were measured by standard plaque assay (PA) on VeroE6 cells obtained from Ralph Baric (referred to as VeroE6_UNC_).

Briefly, 500 µL of serial 10-fold virus dilutions in Opti-MEM were used to infect 4×10^5^ cells seeded the day prior into wells of a 6-well plate. After 1.5 hradsorption, the virus inoculum was removed, and cells were overlayed with DMEM containing 10% FBS with 1.2% microcrystalline cellulose (Avicel). Cells were incubated for 4 days at 33°C, followed by fixation with 7% formaldehyde and crystal violet staining for plaque enumeration. All SARS-CoV-2 experiments were performed in a biosafety level 3 laboratory.

### SARS-CoV-2 neutralization assays

The day before infection, VeroE6/Huh-7.5 cells were seeded at 1×10^4^ cells/well into 96-well plates. Antibodies were serially diluted in DMEM, mixed with target cells and incubated for 60 min at 37°C. Subsequently a constant amount of SARS-CoV-2 was added to achieve 40-50 % virus positive cells. Cells were fixed 18-24 hrafter infection by adding an equal volume of 7% formaldehyde to the wells, followed by permeabilization with 0.1% Triton X-100 for 10 min. After extensive washing, cells were incubated for 1-2 hrat room temperature with blocking solution of 5% goat serum in PBS (005–000-121; Jackson ImmunoResearch). A rabbit polyclonal anti-SARS-CoV-2 nucleocapsid antibody (GTX135357; GeneTex) was added to the cells at 1:1,000 dilution in blocking solution and incubated at 4°C overnight. A goat anti-rabbit AlexaFluor 594 (A-11012; Life Technologies) at a dilution of 1:2,500 was used as a secondary antibody. Nuclei were stained with Hoechst 33342 (62249; Thermo Scientific) at a concentration of 1 μg/mL. Images were acquired with a fluorescence microscope and analyzed using ImageXpress Micro XLS and MetaXpress software (Molecular Devices).

### MAb binding measurements using surface plasmon resonance

Surface Plasmon Resonance (SPR) experiments were performed using a Biacore 8K instrument (GE Healthcare). Human mAbs 2G7A1, 05B04, 05H02 and hybrid mAb 05B04 LC/05D06 HC were captured with a Series S Sensor ship Protein G (Cytiva) at the concentration of 20 nM at the flow rate of 10 μl/min for 60 s. Flow cell one was kept empty and used as a negative control. A concentration series of His-tagged ACE2 1-740aa proteins (4-fold dilutions from a maximum concentration of 500 nM) was injected at 30 µl/min for 240 s followed by a dissociation phase of 2400 s at a flow rate of 30 µl/min. Binding reactions were allowed to reach equilibrium and KD values were calculated from the ratio of association and dissociation constants (KD = kd/ka), which were derived using a 1:1 binding model that was globally fit to all curves in a data set. Flow cells were regenerated with 10 mM glycine pH 1.5 at a flow rate of 30 μl/min for 30 s.

### Spike hACE2 binding and binding inhibition assay

To evaluate whether anti-ACE2 mAbs inhibit spike-ACE2 interaction, 400 ng of each mAb (2G7A1, 05B04, 05H02, or 05B04LC/05D06HC) and a two-fold serial dilution thereof over seven dilutions were mixed with 100 ng of His-tagged ACE2 1-740aa proteins in PBS containing 2% BSA. After 1 hr incubation at 4°C, the mixture was incubated with 10 ng of Wuhan-1 or Omicron S-6P-NanoLuc proteins for 1 hr at 4°C. Then 1 μl of Dynabeads™ His-Tag Isolation and Pulldown magnetic beads (ThermoFisher Scientific) were added into each well. After 1 hr incubation at 4°C, the beads were washed three times and bound NanoLuc activity measured using Nano-Glo Luciferase Assay System and a GloMax® Navigator Microplate Luminometer (Promega).

### Cryo-EM sample preparation, data collection, and structure refinement

Purified 05B04 Fab was mixed with monomeric soluble hACE2 (residues 1-614) at a equimolar concentration for 1h at room temperature. Fab-ACE2 complex was concentrated to 4 mg/mL and deposited on a freshly glow discharged 300 mesh, R1.2/1.3 Quantifoil grid (Electon microscopy sciences). Samples were vitrified in 100% liquid ethane using a Mark IV Vitrobot (Thermo Fisher) after blotting at room temperature and 100% humidity for 3s with Grade 595 filter paper (Ted Pella).

Single-particle cryo-EM data were collected on a Titan Krios transmission electron microscope equipped with a Gatan K3 direct detector, operating at 300 kV and controlled using SerialEm automated data collection software (Mastronarde, 2005). A total dose of 60 e^-^/Å^2^ was accumulated on each movie comprising 40 frames with a pixel size of 0.867Å and a defocus range of −1.0 to −2.6 µm. Data processing was carried out as previously described (Wang *et al*., 2022) using cryoSPARC v3.2 (Punjani et al., 2017) and summarized in Table S3. The global map resolution for the locally-refined reconstruction was 3.3 Å as calculated using the gold-standard Fourier shell correlation of 0.143 criterion.

### Structure modeling, refinement and analysis

Coordinates for initial complexes were generated by docking individual chains from reference structures into cryo-EM density using UCSF Chimera (Goddard et al., 2007). An initial model for the 05B04 Fab – ACE2 structure was generated from coordinates of PDB 7S0B (chain A: 05B04 heavy chain), PDB 7DPM (chain B: 05B04 light chain), and PDB 6VW1 (chain A: hACE2). Models were refined using one round of rigid body refinement with morphing followed by real space refinement in Phenix (Adams et al., 2010). Sequence-updated models were built manually in Coot (Emsley et al., 2010) and then refined using iterative rounds of refinement in Coot and Phenix. Glycans were modeled at potential N-linked glycosylation sites (PNGSs) in Coot. Validation of model coordinates was performed using MolProbity (Chen et al., 2010). Structure figures were made with UCSF ChimeraX (Goddard et al., 2018). Local resolution maps were calculated using cryoSPARC v3.2. Buried surface areas were calculated using PDBePISA (Krissinel and Henrick, 2007) and a 1.4Å probe. Potential hydrogen bonds were assigned as interactions that were <4.0Å and with A-D-H angle >90°. Potential van der Waals interactions between atoms were assigned as interactions that were <4.0Å. Hydrogen bond and van der Waals interaction assignments are tentative due to resolution limitations. ACE2 epitope residues were defined as residues containing atom(s) within 4.5Å of a 05B04 Fab atom.

### hACE2 enzymatic activity assay

To measure the effect of anti-hACE2 mAbs on the catalytic activity of ACE2, various concentrations of 2G7A1, 05B04, 05H02, and 05B04LC/05D06HC (2 μg/ml, 10 μg/ml, 50 μg/ml) were mixed with ACE2 at 0.2 μg /ml. ACE2 enzymatic activity was then measured using the ACE2 Inhibitor Screening Assay Kit (BPS Bioscience) following the manufacturer’s instructions. The intensity of the fluorescent product of the ACE2 reaction product was detected at 555nm/585 nm (excitation/emission) with Clariostar Plus Microplate Reader (BMG Labtech). MLN-4760 (Sigma, #5306160001) served as a positive control ACE2 inhibitor.

### Flow cytometric analysis of cell surface hACE2 binding by human anti-hACE2 mAbs

To evaluate the ability of anti-ACE2 mAbs to bind to cell surface ACE2 by flow cytometry, A549 cells (human alveolar basal epithelial cells) were engineered to express hACE2. The hACE2 or macaque (mac)ACE2 expressing cells were detached from plates with 10 mM EDTA in PBS and 10^5^ cells incubated in the absence or the presence of human anti-ACE2 mAbs (2 μg/ml) for 2 hr at 4°C. After washing, the cells were incubated with AlexaFluor™ 488 conjugated goat anti-human IgG (ThermoFisher Scientific). Flow cytometry was performed using Attune® NxT Acoustic Focusing Cytometer (ThermoFisher Scientific). The same procedure was applied to parental, unmodified A549 cells as a negative control for nonspecific cell surface binding.

### hACE2 internalization assay

To determine whether the anti-ACE2 mAbs induced ACE2 internalization, live A549 cells expressing hACE2 receptor with an HA-epitope tag appended to its intracellular C-terminus were incubated with anti-human ACE2 mAbs (1 μg/ml) for 1 hr at 37°C. Then, cells were fixed with 4% PFA/PBS, treated with 10 mM glycine, and permeabilized with 0.1% Triton X-100. Total human ACE2-HA was then detected with mouse anti-HA.11 antibody (1 μg/ml). The internalization of the ACE2-HA protein and anti-hACE2 mAbs was then evaluated by staining with goat anti-mouse Alexa Fluor™ 594 (to detect the HA-tagged hACE2) and/or goat anti-human Alexa Fluor™ 488 antibodies (to detect the hACE2 mAbs) (ThermoFisher Scientific). Images were captured using a DeltaVision OMX SR imaging system (GE Healthcare).

### Analysis of anti-ACE2 mAbs pharmacokinetics in mice

Six week old hACE2-knock-in female mice, in which human ACE2 cDNA replaces the endogenous mouse ACE2 sequences, were obtained from Jackson Labs (B6.129S2(Cg)-Ace2tm1(ACE2)Dwnt/J, strain 035000). After acclimatization for 2 weeks, these mice received subcutaneous injections of 250 μg of human anti-hACE2 mAbs per mouse (n=5). Mice were bled on day 0, day 1, day 3, day 7, and day 14 with blood collected into Microvette® CB 300 Serum (Sarstedt).

Serially diluted mouse plasma (five-fold serial dilution over four dilutions from a maximum volume of 0.5 μl) was diluted in PBS buffer containing 2% BSA and mixed with 30 ng of His-tagged ACE2(1-740aa)-NanoLuc protein. After 1 hr incubation at 4°C, the mixture was incubated with 3 μl of Dynabeads™ Protein G magnetic beads (ThermoFisher Scientific). After 1 hr rotation at 4°C, the beads were washed three times and bound NanoLuc activity was measured using Nano-Glo Luciferase Assay System and a GloMax® Navigator Microplate Luminometer (Promega). To construct standard calibration curves for measurement of mAb levels in plasma, 100 ng of mAbs (2G7A1, 05B04, 05H02, or 05B04LC/05D06HC) were five-fold serially over seven dilutions were and mixed with ACE2(1-740aa)-NanoLuc proteins. MAb:ACE2(1-740aa)-NanoLuc complexes were captured and quantified in parallel with those formed using the plasma samples from mAb infused mice.

### SARS-CoV-2 challenge experiments in hACE expressing mice

Six week old hACE2-knock-in female mice, in which human ACE2 cDNA replaces the endogenous mouse ACE2 sequences, were obtained from Jackson Labs (B6.129S2(Cg)-Ace2tm1(ACE2)Dwnt/J, strain 035000). After acclimatization for 2 weeks, the mice (five mice per treatment group) were injected retro-orbitally with 250 μg (equivalent to ∼12.5 mg/kg) of anti-hACE2 mAbs. At 2 days after mAb injection, mice were challenged intranasally with SARS-CoV-2, USA_WA/2020 P3, 2×10^5^ PFU/mouse (virus titers determined on VeroE6 cells). At 3 days after infection, mouse lungs were dissected and homogenized in TRIzol. Chloroform was added to induce phase separation. Then after centrifugation, RNA in the aqueous phase was precipitated with isopropanol and, after wash with ice-cold 75% ethanol, dissolved in nuclease-free water. The number of viral genomes per microgram of total lung RNA was measured by qRT-PCR, using Power SYBR Green RNA-to-CT 1-Step Kit (ThermoFisher Scientific) on StepOne Plus Real-Time PCR system (Applied Biosystems). The primers used target RNA sequences encoding the nucleocapsid protein: 2019-nCoV_N1-F: 5′-GACCCCAAAATCAGCGAAAT-3′ and 2019-nCoV_N1-R: 5′-TCTGGTTACTGCCAGTTGAATCTG-3′. The standard was obtained from Integrated DNA technologies (2019-nCoV_N_Positive Control 10006625)

## Supplementary information

**Table S1:**
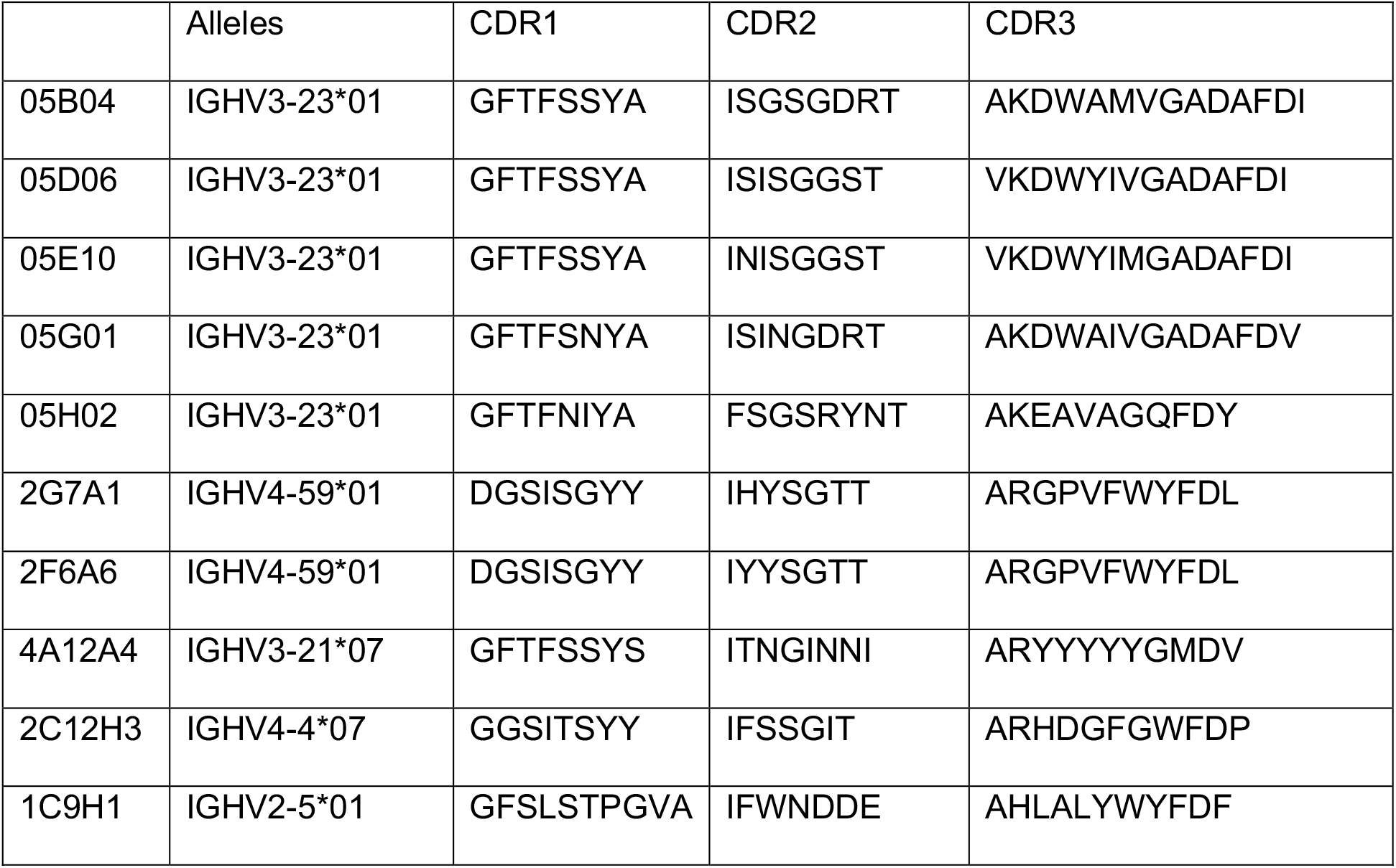
CDR sequences of anti-ACE2 mAb heavy chains.

**Table S2:**
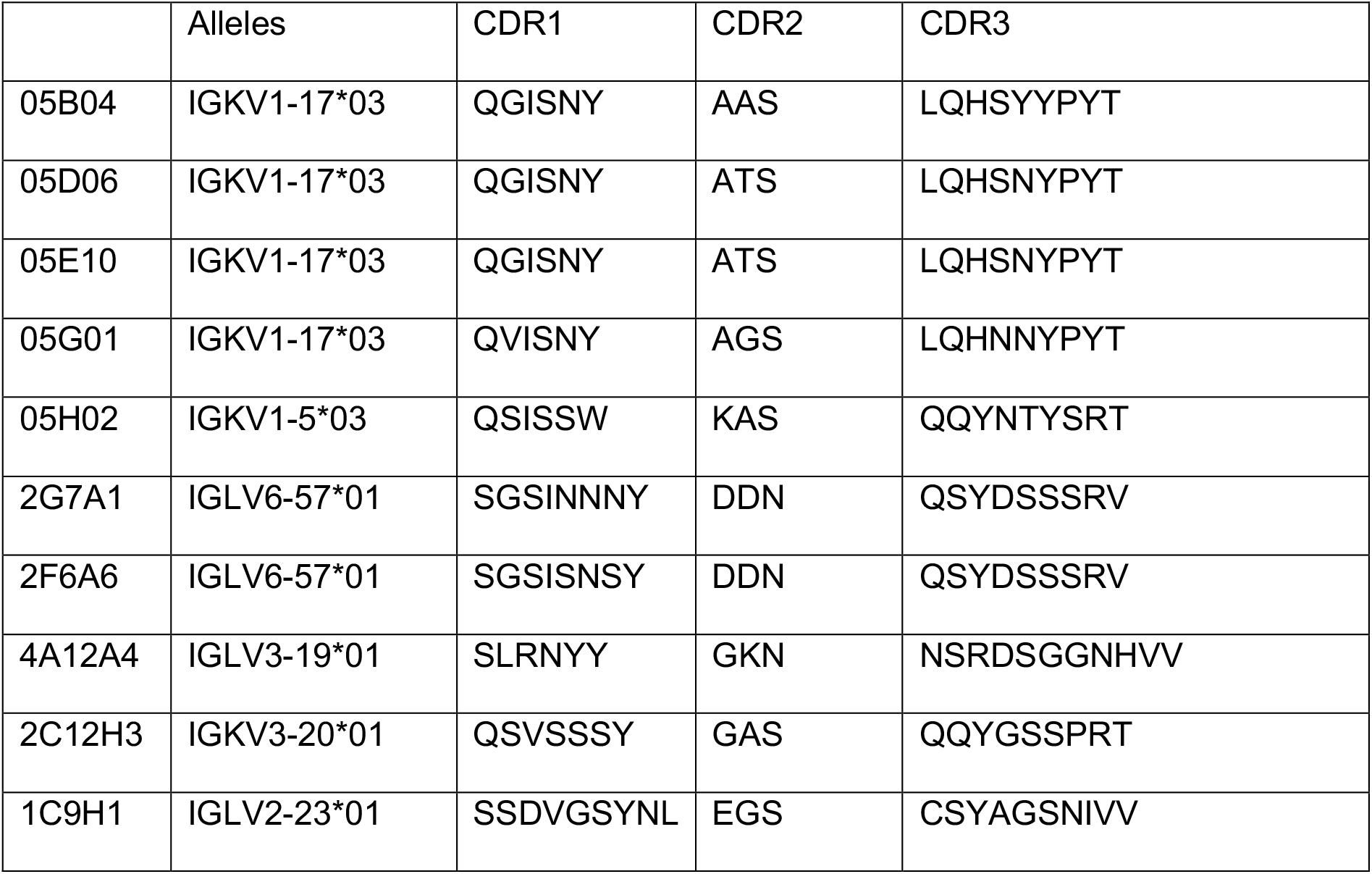
CDR sequences of anti-ACE2 mAb light chains.

**Table S3.** Cryo-EM data collection, refinement and validation statistics.

|  |  |
| --- | --- |
|  | **05B04 Fab**<br>**hACE2<sub>1-614</sub>**<br>**complex** |
| **PDB** | **8E7M** |
| **EMD** | **27939** |
| **Data collection conditions** |  |
| Microscope | Titan Krios |
| Camera | Gatan K3 Summit |
| Magnification | 130,000x |
| Voltage (kV) | 300 |
| Dose rate (e<sup>-</sup>/pixel/s) | 17.6 |
| Electron dose (e<sup>-</sup>/Å<sup>2</sup>) | 60 |
| Defocus range (μm) | 1.0-2.6 |
| Pixel size (Å) | 0.8677 |
| Micrographs collected | 6,599 |
| Micrographs used | 4,907 |
| Total extracted particles | 2,055,266 |
| Final refined particles | 490,485 |
| Symmetry imposed | C1 |
| Nominal Map Resolution (Å) |  |
| FSC 0.143 (unmasked/masked) | 3.9/3.5 |
| **Refinement and Validation** |  |
| Initial model | 6VW1, 7S0B, 7DPM |
| Model Resolution (Å) |  |
| FSC 0.143 | 3.3 |
| Number of atoms |  |
| Protein | 8,028 |
| Ligand | 146 |
| MapCC (global/local) | 0.79/0.75 |
| Map sharpening B-factor | 112.6 |
| R.m.s. deviations |  |
| Bond lengths (Å) | 0.004 |
| Bond angles (°) | 0.814 |
| MolProbity score | 1.74 |
| Clashscore (all atom) | 7.89 |
| Poor rotamers (%) | 0 |
| Ramachandran plot |  |
| Favored (%) | 95.5 |
| Allowed (%) | 4.5 |
| Disallowed (%) | 0 |

**Figure S1.**
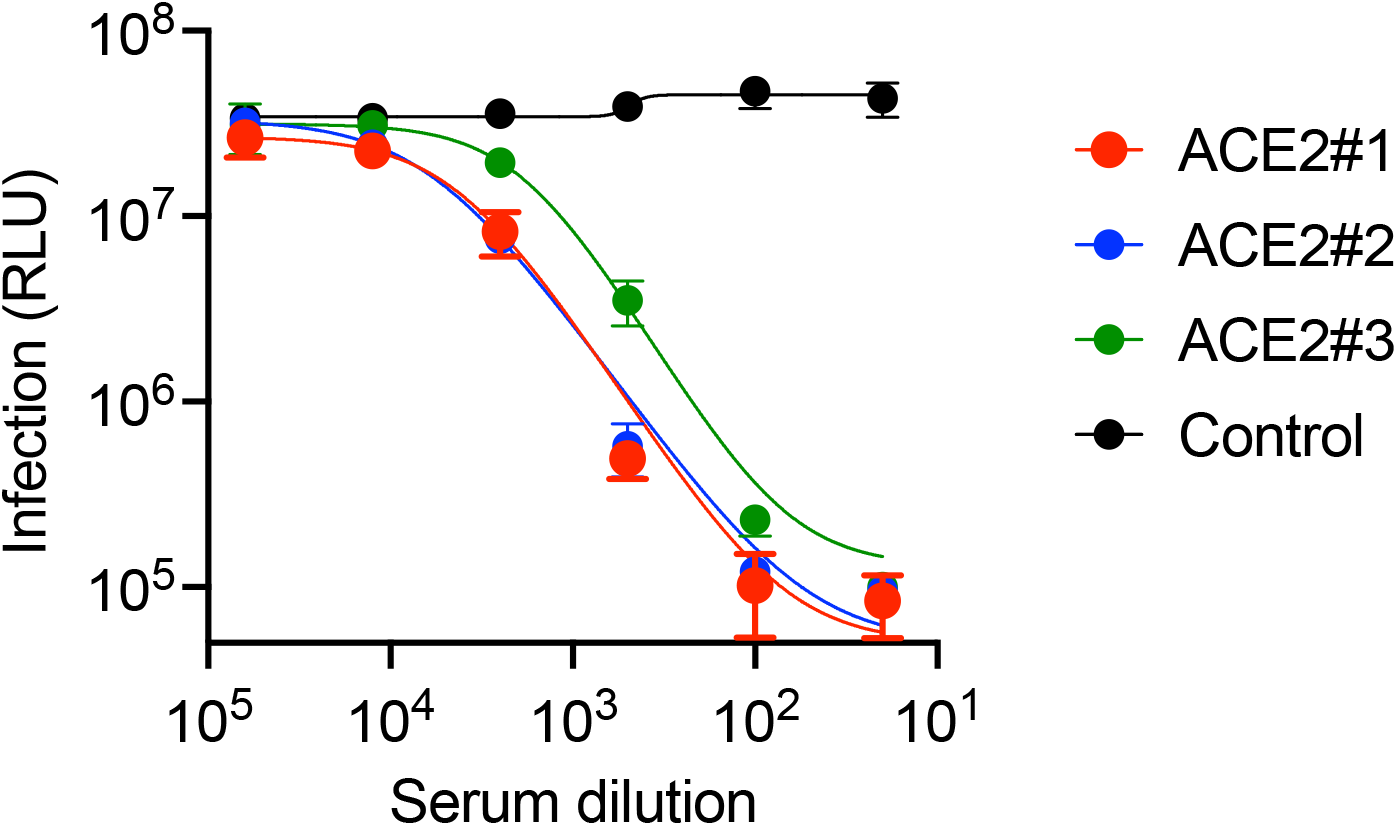
Inhibition of HIV-1 based pseudotyped virus infection by anti-hACE2 mAbs mouse serum. Heat inactivated serum from hACE2-8XHis immunized mice or an unimmunized control was incubated with Huh-7.5 target cells. Cells were then infected SARS-CoV-2 pseudotyped HIV-1 and infection was measured using NanoLuc luciferase assay 48h later.

**Figure S2.**
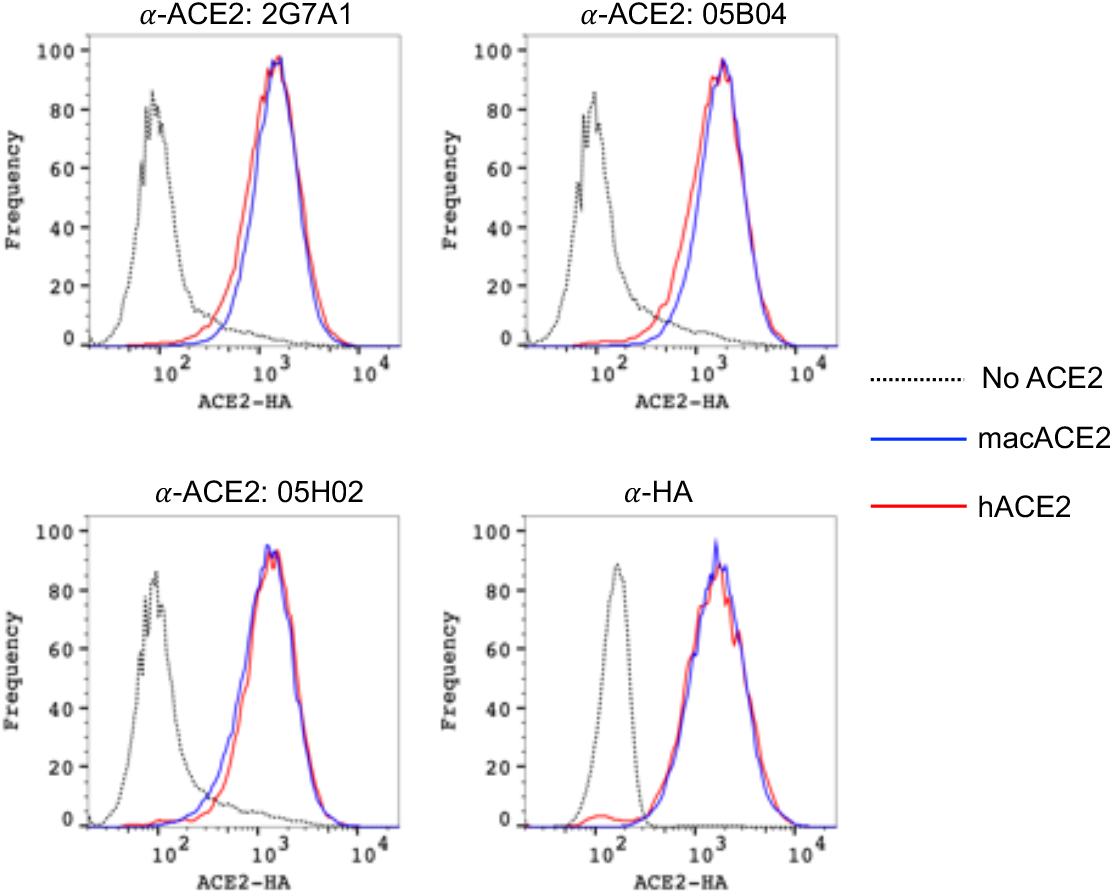
Binding of anti-hACE2 antibodies to human and macaque ACE2. Parental A549 cells (dashed lines), or A549 cells stably expressing HA-tagged macaque ACE2 (blue, macACE2), or HA-tagged human ACE2 (red, hACE2) were incubated in the presence of the indicated human anti-hACE2 antibodies or anti-HA antibody. The cells were then incubated with Alexa Fluor 488 conjugated goat anti-human IgG (for anti-human ACE2 antibodies) or Alexa Fluor 488 goat anti-mouse IgG (for anti-HA antibody) and then analyzed by flow cytometry.

**Figure S3.**
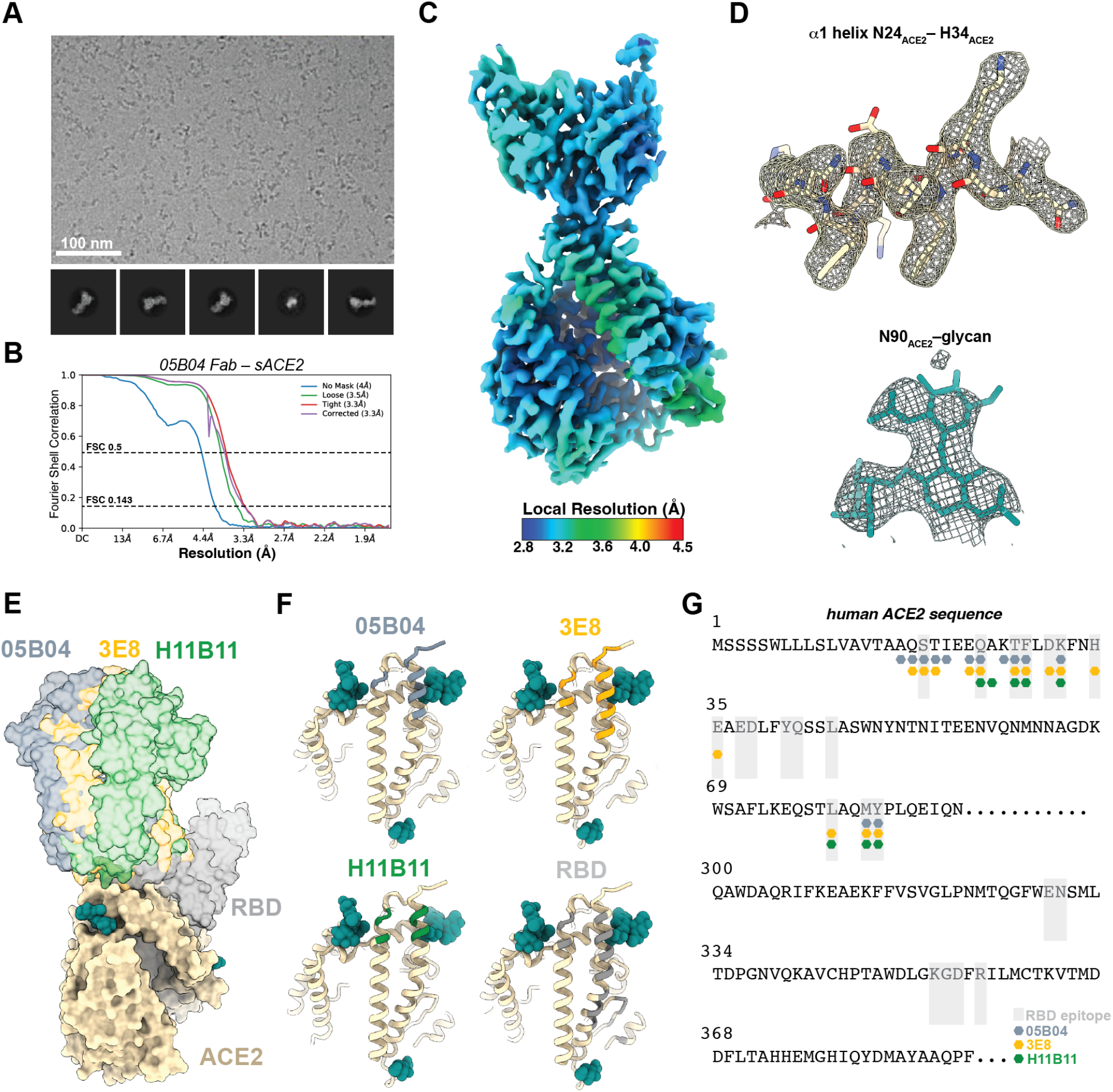
Cryo-EM data processing, validation, and structure comparison. (A) Representative micrograph and 2D class averages for 05B04-ACE2 data collection. (B) Gold-standard FSC plot calculated in cryoSPARC for local refinement. (C) Local resolution estimate for 05B04-ACE2 locally-refined map. (D) Representative cryo-EM density contoured at 7σ for select regions. (E) Structural superposition of anti-ACE2 mAbs 05B04 (slate blue), 3E8 (orange), h11b11 (green) and SARS-CoV-2 RBD (gray) modeled on sACE2 (wheat). (F,G) Comparison of anti-ACE2 mAb epitopes visualized on an ACE2 model (F) or highlighted on the ACE2 sequence (G). anti-ACE2 mAb epitopes are colored slate blue, orange, and green for 05B04, 3E8, and h11b11, respectively. The SARS-CoV-2 RBD epitope is denoted as shaded gray boxes (G).

**Figure S4.**
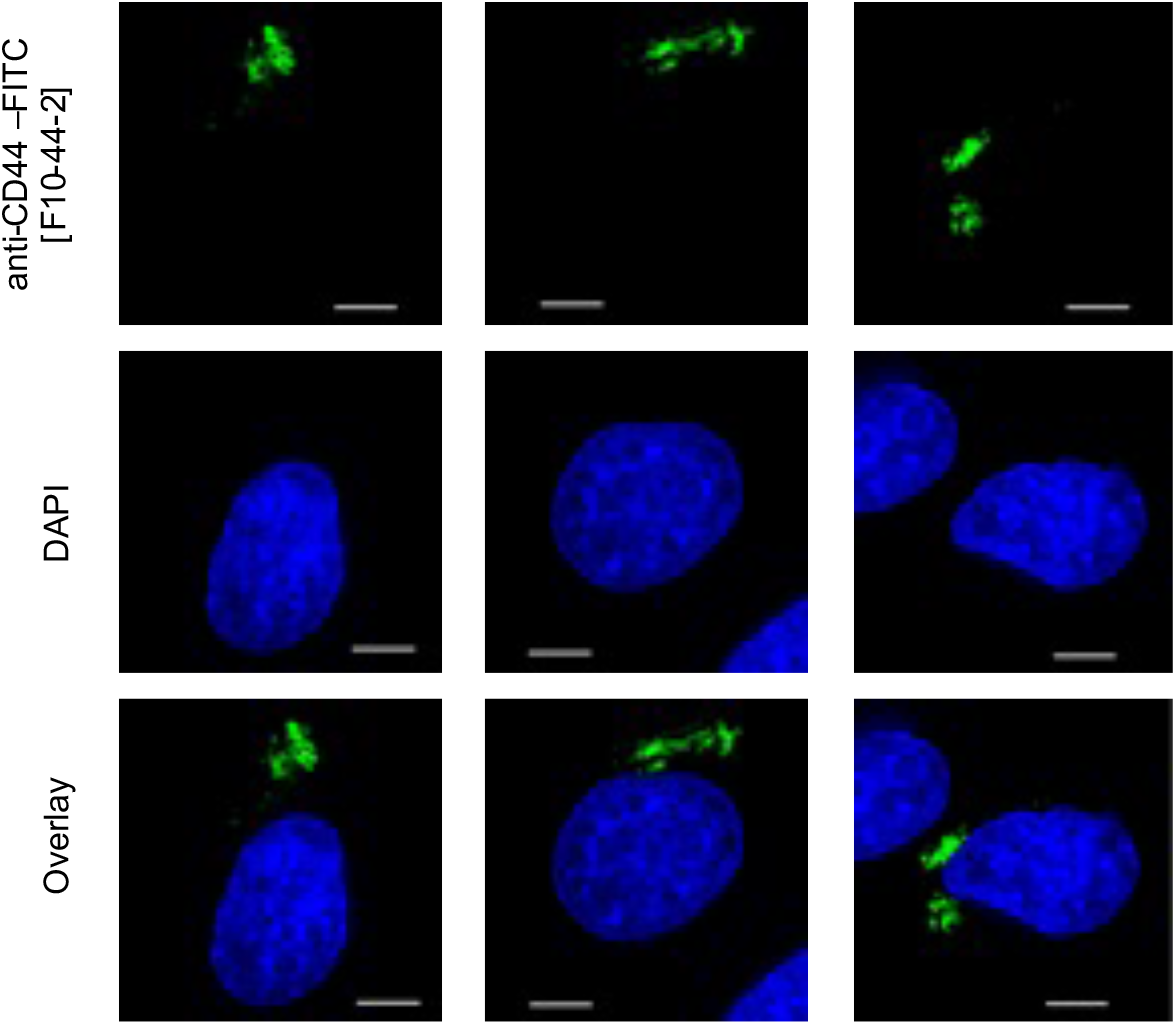
Internalization of a control anti-CD44 antibody in A549/ACE2-HA cells. Localization of anti-CD44 antibody (green, upper panels) following incubation of live A549/ACE2-HA cells with FITC-labelled anti-CD44 (F10-44-2). Three representative fields are shown. Blue stain (DAPI) indicates cell nuclei.

**Figure S5.**
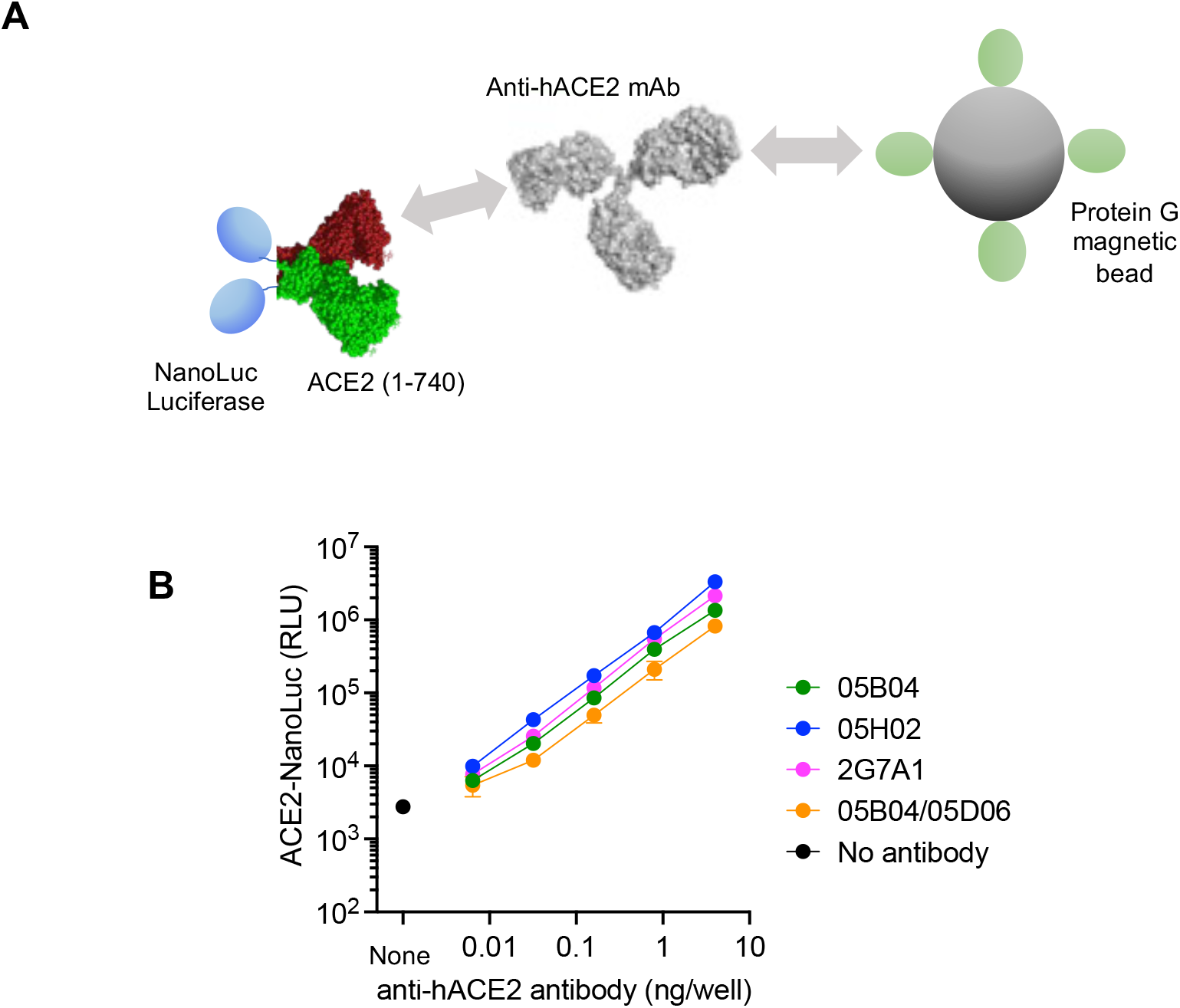
Determination of anti-hACE2 antibody levels using ACE2-NanoLuc fusion protein. (A) Schematic representation of the NanoLuc assay in which NanoLuc luciferase was appended to the C-termini of the hACE2 extracellular domain (1-740 aa). The fusion protein was incubated with anti-ACE2 mAbs and complexes were then captured using Dynabeads™ Protein G magnetic beads. (B) For the standard curves, 40 ng of the indicated anti-ACE2 mAbs and five-fold serial dilutions thereof, were mixed with 30 ng of ACE2(1-740 aa)-NanoLuc protein. After 1 hr incubation at 4°C, the mixture was incubated with Dynabeads™ Protein G magnetic beads. The beads were washed and bound NanoLuc activity was measured.

